# Much higher covariation with foveation timing by superior colliculus than primary visual cortical neuronal activity

**DOI:** 10.1101/2025.07.10.664256

**Authors:** Carlotta Trottenberg, Yue Yu, Tong Zhang, Matthias P. Baumann, Tatiana Malevich, Shweta Prasad, Ziad M. Hafed

**Affiliations:** Werner Reichardt Centre for Integrative Neuroscience, University of Tübingen, Tübingen, Germany; Hertie Institute for Clinical Brain Research, University of Tübingen, Tübingen, Germany

**Keywords:** Superior colliculus, primary visual cortex, saccades, eye movements, saccadic reaction time, active vision

## Abstract

The superior colliculus (SC) is a potent driver for orienting movements, but it also possesses short-latency visual responses. Even though such responses largely derive from direct primary visual cortical (V1) input, their relationship to V1 activity has never been analyzed with the same experimental subjects and stimuli. Here, we revisited pioneering observations that trial-by-trial variability in either SC or V1 visual responses may predict eye movement timing variability. We found that V1 covariation with behavioral variability was virtually non-existent in comparison to the SC, whether with respect to visual response onset latency, visual response strength, or pre-stimulus state. By far, our largest predictor of behavioral variability was visual response strength in SC visual-motor neurons. These results suggest that the SC reformats its sensory inputs for exploitation of the SC’s proximity to the motor periphery; V1 aids in jumpstarting the sensing process, but the SC much more directly supports visually-driven orienting.

## Introduction

Visually-driven orienting movements are a fundamental component of active sensing in many organisms. In primates, rapid foveating saccadic eye movements are often thought of as a “visual grasp reflex” ^1–4^, given how effectively eccentric visual stimulus events can attract such eye movements. However, and perhaps remarkably in the case of a single visual onset in an otherwise uniform visual field, saccades towards the very same appearing stimulus can exhibit large variability in their reaction times ^5–9^. One aspect of such variability is related to the asynchronous nature of exogenous stimuli relative to the internal oculomotor rhythm ^10^, and others include cognitive processes such as target selection ^9,11^ and decision making ^9,12^. Yet others relate to motor command equilibria ^13^. However, none of these processes can even begin before sensory detection of stimulus occurrence takes place; thus, variability in the sensing process itself can be an additional contributor to eye movement timing variability.

Previous pioneering work has demonstrated that visual stimulus properties, such as intensity, are indeed reflected in both the average visual response latencies of superior colliculus (SC) neurons as well as in the subsequent average saccadic reaction times ^14^. Later studies have also shown how both SC visual response strength and visual response timing can reliably predict average saccadic reaction times ^15,16^, and how this could also happen for different visual feature dimensions ^17–19^ as well as when the same responses to the same stimuli are modified by processes such as saccadic suppression ^19–21^. In the latter case, this indicates that trial-by-trial variability in SC visual responses (to the same stimulus) can correlate with trial-by-trial saccadic reaction time variability, which has also been demonstrated ^15^.

All of the above results support the view that SC visual responses are integral to saccade generation ^22^, but they also raise a fundamental question: what is special about these visual responses? For example, in equally important pioneering work, this time in the primary visual cortex (V1) ^23^, it was shown that variability in V1 visual response timing to repeated presentations of the same visual stimulus also correlates well with the timing of saccadic eye movements. Given that the majority of visual inputs to the SC anatomically derive from a direct connection originating in V1 ^24–30^, does this mean that the SC’s contributions to behavioral variability are merely inherited from V1? Answering this important question requires measuring SC and V1 neuronal variability in the very same experimental subjects and with the very same stimuli and behaviors, which we did here.

We measured trial-by-trial variability in visually-guided saccade timing, and we related it to trial-by-trial variability in visual response onset latency, visual response strength, and pre-stimulus neuronal activity. Visual response onset latency allowed us to assess the importance of a temporal code in dictating the timing of saccadic eye movements ^23^, whereas visual response strength revealed the importance of sensory evidence integration. Of course, in the case of the SC, it has been known for around five decades or more already that saccade-related neuronal activity (so-called “motor bursts”) essentially perfectly predicts saccade onset times ^31,32^. However, our purpose here was not to revisit this highly established fact; instead, we aimed to document a much-needed direct comparison between the influences of SC and V1 stimulus-related activity on behavioral variability. Even though the primate SC receives the majority of its inputs from V1 ^24,25,27–29,33^, we found a striking difference in how its visual responses correlated to foveating eye movement timing, with the V1 effects being effectively non-existent when compared to the SC. We also found that a rate, rather than temporal, code dominated the relationship between trial-by-trial SC visual response variability and saccadic reaction times. Relying on such a rate code, necessitating temporal integration, can reduce sensitivity to noise, which can erroneously trigger movements through the SC’s strong excitatory downstream projections.

## Results

We collected neuronal activity from the SC and V1 (Fig. 1A) while rhesus macaque monkeys performed a saccade towards the location of a suddenly appearing eccentric visual stimulus in the recorded neurons’ response fields (RF’s) (Fig. 1B). The stimulus appeared simultaneously with the offset of the initial fixation target (Fig. 1C), and it consisted of a luminance disc of 0.51 deg radius; across trials, the stimulus could have different levels of Weber contrast or luminance polarity (bright or dark) relative to the surrounding gray background (Methods) (Fig. 1D) ^18^. We focused on trial-by-trial variability in the onset timing of the foveating saccadic eye movements, and we asked what aspects of stimulus-aligned neuronal activity in the two brain areas could better predict such variability, and to what extent.

**Figure 1.**
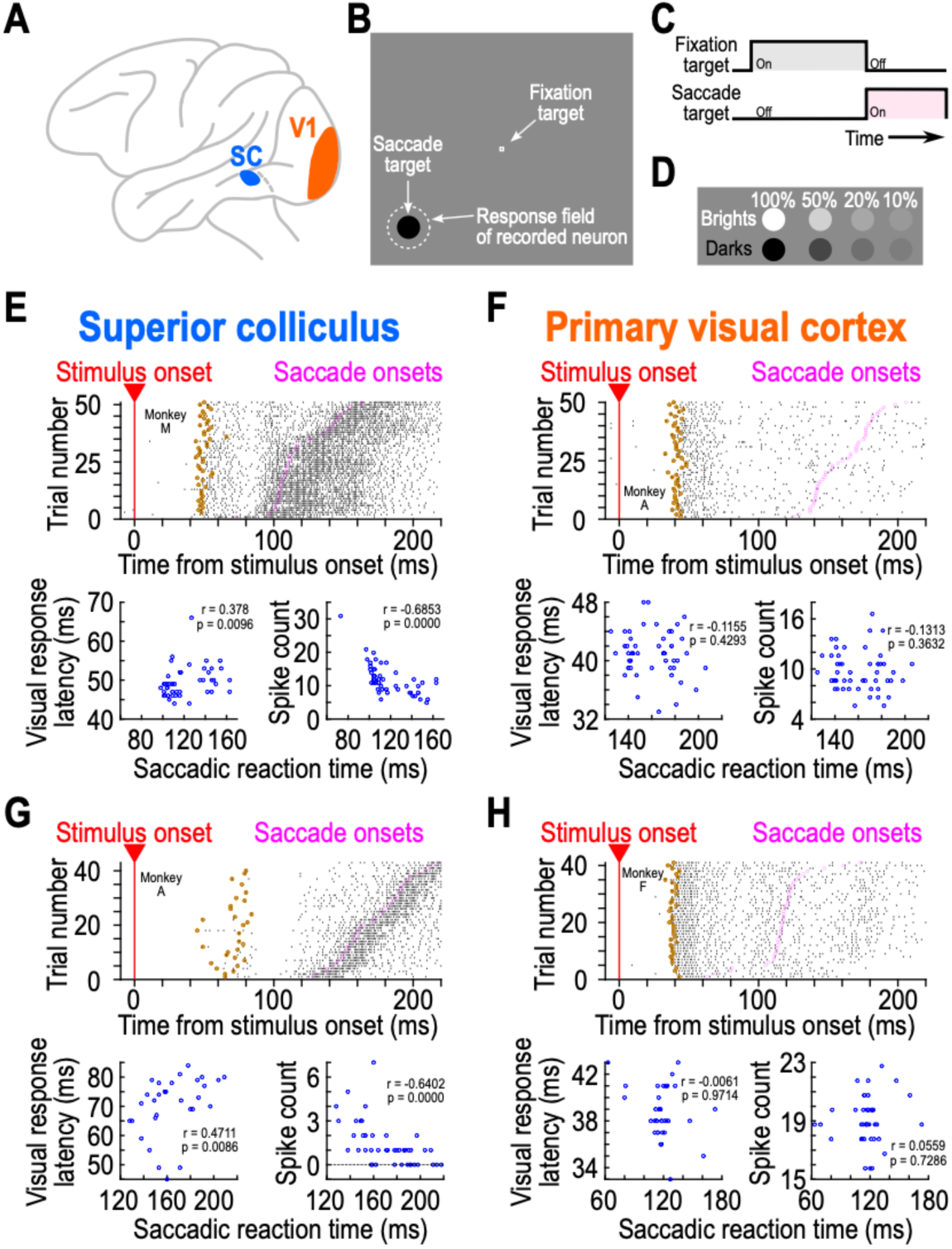
Qualitatively similar, but quantitatively different, visual responses in the superior colliculus (SC) and primary visual cortex (V1). **(A)** We compared visual responses in the SC (blue) and V1 (orange). **(B)** The task was a visually-guided saccade task, and the saccade target appeared within the recorded neurons’ response fields (RF’s). **(C)** Timeline of events in each trial. The appearance of a fixation spot initiated a trial, after some delay (Methods), the saccade target appeared, coincident with the disappearance of the fixation spot. **(D)** The saccade targets could have different luminance polarities (dark or bright) and contrasts. **(E)** The top panel shows neuronal activity of an example SC neuron when the monkey oriented towards a suddenly appearing eccentric stimulus (50% dark contrast). Trials are sorted by the time of saccade onset (magenta; saccade onsets). The neuron emitted a short-latency visual burst shortly after stimulus onset in its RF; brown indicates the estimated visual response onset time on each trial (Methods). On the slow saccadic reaction time trials, there were later visual response onsets (compare the timing of the brown symbols across trials); additionally, the visual responses were weaker (there were fewer spike counts in the visual bursts for the slow trials). The bottom panels relate the visual response latency (left) and spike count in the visual burst interval (right) to the time of saccade onset. There were both later and weaker visual bursts in the slow reaction time trials (Spearman correlation coefficient values shown within each panel). **(F)** For an example V1 neuron, this relationship was not present. Note how the SC neuron (**E**) also exhibited a saccade-related burst at eye movement onset, whereas the V1 neuron did not. **(G, H)** An additional pair of example neurons from the two brain areas. Once again, visual responses in the SC neuron were later and weaker for slow than fast saccadic reaction time trials, but this was not systematically the case for the V1 neuron. Also see Fig. S2 for eight additional example neuron results (under the same stimulus conditions), and Fig. 3 for across-trial firing rate averages of the neurons.

### Superior colliculus visual response latencies are better predictors of saccadic reaction times than primary visual cortical visual response latencies

Figure 1E (top) shows the activity of an example SC neuron (from monkey M) when the appearing disc was darker than the surrounding background and had a 50% Weber contrast. The top panel shows spike rasters, indicating that the neuron emitted a visual response shortly after stimulus onset. The neuron, being a visual-motor neuron, also emitted a subsequent saccade-related motor burst, which (as expected ^31,32^) was time-locked to saccade onset. To estimate the neuron’s visual response onset latency from individual trials, we convolved the within-trial spike times with an asymmetric firing rate kernel that avoided the blurring of response onset estimates backwards in time (Methods; Fig. S1) ^23^. We then used the individual-trial firing rate estimates to compare post- to pre-stimulus activity and infer the visual response onset time (Methods) ^23^. For the shown example neuron, the estimated onsets of visual responses on individual trials are shown by the brown circles. We also detected the onsets of foveating saccades (magenta circles in the figure; Methods), and we sorted trials according to these saccades’ onset times. This example neuron’s visual response onset was delayed on the trials in which the subsequent saccade was late (larger trial numbers in the top panel of Fig. 1E), and this effect can also be seen in the bottom left panel of Fig. 1E. In this panel, we plotted visual response onset latency (the brown circles in the top panel) against saccadic reaction time (the magenta circles). The Spearman correlation coefficient between trial-by-trial visual response onset latency and saccadic reaction time was 0.378 (p=0.0096). In contrast, for an example V1 neuron (Fig. 1F), there was no positive correlation between trial-by-trial visual response timing and trial-by-trial saccade timing; the Spearman correlation coefficient in this case was -0.1155 (p=0.4293). Similar conclusions could also be reached from two additional example neurons, one from the SC (Fig. 1G) and one from V1 (Fig. 1H), recorded from two additional monkeys (0.4711 versus -0.0061 Spearman correlation coefficients, respectively, indicating better covariation with behavior in the SC than in the V1 neuron; p=0.0086 and p=0.9714 for the SC and V1 neurons, respectively). Thus, at least for these four example neurons, there was a clearer positive correlation between visual response onset latency and saccadic reaction time in the SC than in V1. Figure S2 shows eight additional example neurons (four from the SC and four from V1) exhibiting similar results.

Across neurons in either the SC or V1, we first measured, for each neuron, the Spearman correlation coefficient between trial-by-trial visual response onset latency and trial-by-trial saccadic reaction time. We then obtained summary statistics across the population. In the SC, there was a clear positive correlation coefficient value for all tested contrasts, consistent with earlier reports ^15,16^. For example, for the dark contrasts shown in Fig. 2A, the average correlation coefficient was around 0.1 for 10-20% contrast levels, and around 0.06 for the higher contrasts (Fig. 2A; results for bright contrasts were qualitatively similar and will be more explicitly discussed below). Critically, these values were greater than approximately 2-6 times larger than the values observed in V1 for the very same visual stimuli (Fig. 2A). That is, across the population, SC visual response onset latencies were a markedly better predictor of saccade timing than V1 visual response onset latencies. This was also not due to our stimuli being somehow suboptimal for the V1 neurons; to the contrary, we systematically observed strong V1 visual responses to these stimuli (Fig. 3).

**Figure 2.**
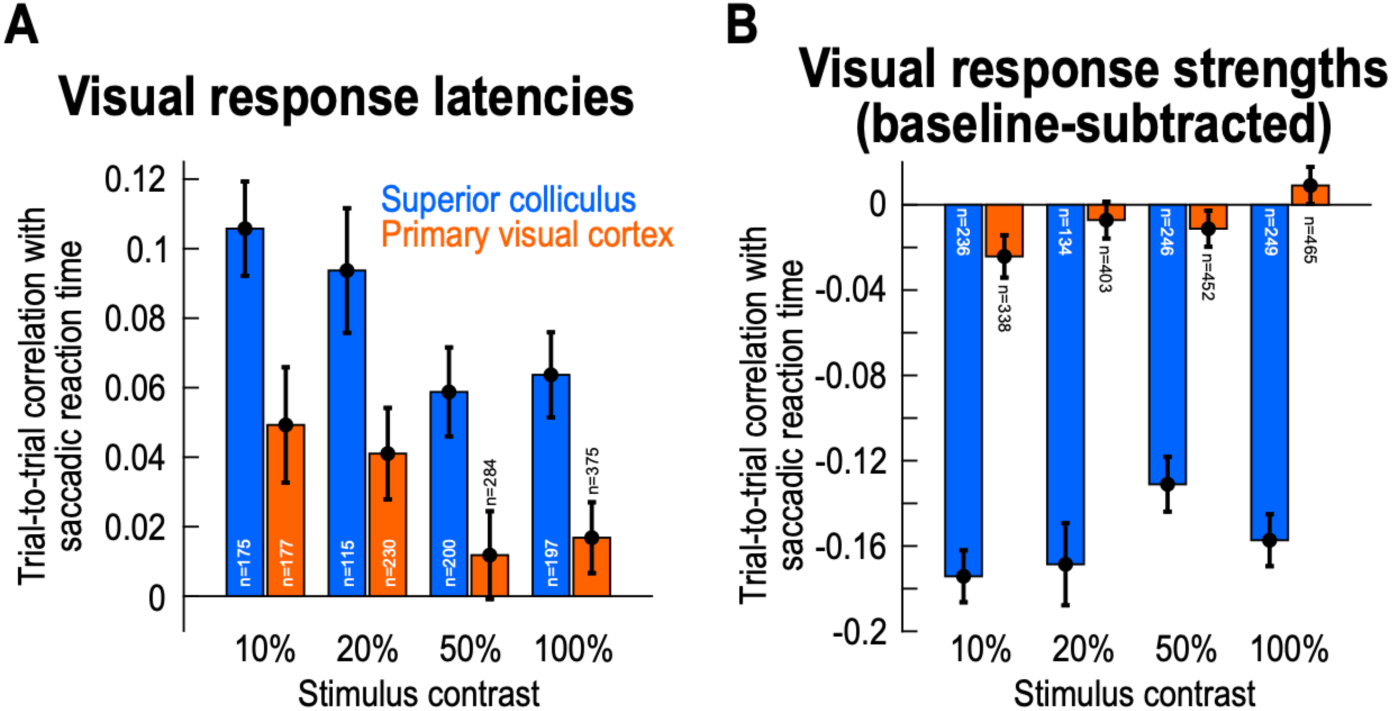
Much higher trial-by-trial correlations with saccade timing in the SC than in V1, for both visual response latencies as well as visual response strengths. **(A)** For each stimulus Weber contrast that we tested (all being darker than the gray background in this figure; Methods), we calculated the Spearman correlation coefficient value (within any given neuron) between the individual trial saccadic reaction times and the individual trial neuronal visual response latencies. We then summarized such correlation coefficient values across all neurons in our population. Positive correlation coefficients indicate later visual response latencies for longer saccadic reaction times, and earlier visual response latencies for shorter reaction times. The SC correlation coefficients were approximately 2-6 times greater than the V1 correlation coefficients. **(B)** We also characterized the correlations between trial-by-trial visual response strengths and trial-by-trial saccadic reaction times; the former were defined as the spike counts within a visual burst interval, after subtracting the baseline pre-stimulus activity (Methods). Negative correlation coefficients indicate weaker visual burst strengths for later saccades, and stronger visual burst strengths for earlier saccades. The correlation coefficients in V1 were practically zero, but they were more clearly negative in the SC, and even more so, in absolute value, than the visual response latency correlations in **A**. Thus, visual response strength in the SC is significantly better correlated with the timing of saccadic behavior than visual response latency; and, for both measures, the effects are much weaker in V1. Error bars denote SEM across neurons, and the numbers of neurons for each analysis are included in the figure. Note that in **A**, results are only shown for the neurons in which our criteria for sufficient visual response onset latency measurements were met (Methods), explaining the slightly lower neuron counts than in **B**. Also note that for each neuron, multiple contrast levels were tested (Methods). Therefore, the total number of recorded neurons in each brain area (reported in Methods) was not the sum of the shown numbers in each panel. Figure S3 shows individual monkey results, and Fig. S4 shows the raw histograms underlying all of the data in the current figure.

**Figure 3.**
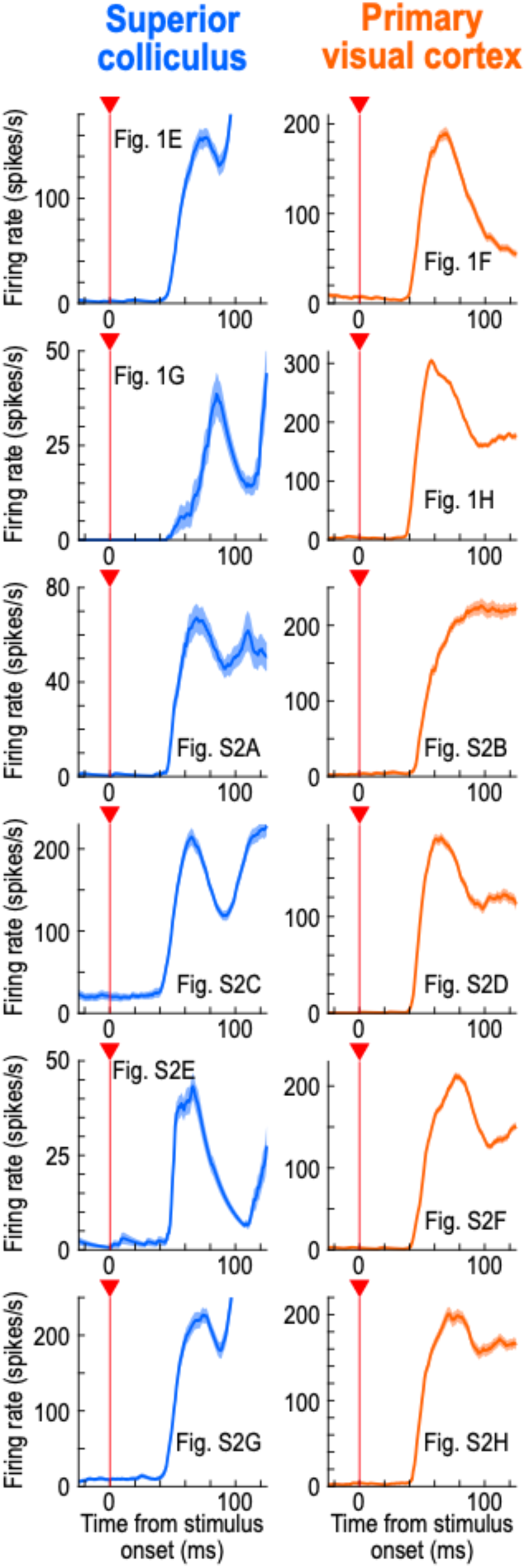
Strong visual responses to our used stimuli by the recorded V1 neurons. This figure shows average firing rates across trials for all of the example neurons shown in Figs. 1, S2 (for the 50% dark contrast). Here, we plotted the neuronal responses in the visual epoch to demonstrate that the V1 results of Fig. 2 were not necessarily explained by our stimuli being somehow suboptimal for V1 neurons. We consistently observed strong V1 visual responses for these stimuli (and we saw even higher V1 firing rates for 100% dark contrasts). Error bars denote SEM across trials, and each panel indicates which figure panel in Figs. 1, S2 the shown firing rate data correspond to. The numbers of trials in each panel were resported in Figs. 1, S2.

Statistically, we performed a Mann-Whitney U test comparing the SC and V1 correlation coefficient values across all contrasts (Methods; Fig. 2A). These correlation coefficients were significantly higher in the SC than in V1 (U=663340, z=5.76, p<0.0001, n_SC_=688, n_V1_=1066, median_SC_=0.0804, median_V1_=0.0217). Moreover, at each contrast level individually, the SC correlation coefficient values were again significantly higher than the V1 correlation coefficient values after Bonferroni correction (Mann-Whitney U tests; p<0.01 in each comparison). Therefore, across the population, we confirmed the existence of a substantially stronger covariation with behavioral variability in the SC’s visual response onset latencies than in V1.

We also noticed a trend for higher correlation coefficient values to behavioral variability at low contrasts, even for V1 (Fig. 2A). For example, at 10% contrast, a Wilcoxon Signed-Rank test against zero revealed significantly positive correlations in the V1 population after Bonferroni correction (W=9791, z=2.80, p=0.0050, n=177). This was also the case at 20% contrast (W=16114, z=2.80, p=0.0051, n=230). Interestingly, when we inspected individual monkey results, we found that this observation was essentially entirely dominated by monkey F’s V1 data (see Fig. S3A, D), and that this same monkey had much faster saccadic reaction times than the other two animals (Fig. S3C, F, I). These observations might suggest that for very fast saccadic reaction times, approaching the reaction times of express saccades ^34^, V1 visual responses may boost the drive of foveating eye movements. The fact that this effect was stronger at low contrasts (Figs. 2A, S3D) might additionally explain why some significantly positive population correlation coefficient values between V1 visual response onset latencies and saccadic reaction times were previously reported ^23^; in that study, generally low contrasts were investigated ^23^. Finally, in the SC, all three individual monkeys showed consistent results (Fig. S3A, D, G), further supporting the idea that SC visual responses are better aligned with behavioral variability than V1 visual responses.

### Superior colliculus visual response strengths are even better predictors of saccade timing

Our results above suggest that the first stimulus-evoked action potential by a neuron is positively correlated with trial-by-trial saccadic reaction time variability in the SC, and that this effect is weaker in V1. However, perceptual decisions aiding in foveation involve not only jumpstarting the sensing process via the timing of the first spike in a sensory response (Fig. 2A), but they also require sensory evidence integration ^18,35–38^. For example, and particularly for weak stimuli, the onset of a neuronal visual response to a stimulus may itself be difficult to assess by recipient neurons. Moreover, given how even individual action potentials by individual SC neurons can very easily modify or even trigger new eye movements ^10,39,40^, even if these action potentials were purely visually-driven, it might be reasonable to hypothesize that for the SC, in particular, sensory evidence integration (a rate code) is more effective than a temporal code (visual response onset latency); this way, the final oculomotor behavior may be much more immune to potential spiking variability in SC neuronal activity than otherwise. To test this, we investigated how visual response strength (the number of stimulus-evoked spikes emitted by an individual neuron during a so-called visual response epoch; Methods) could predict saccadic reaction time in both the SC and V1.

Visual response strength overwhelmingly dominated the predictive powers of visual responses in the SC about saccade timing variability, and the dissonance from V1 effects that we observed in Fig. 2A was also rendered much more substantial. Consider, first, the same example SC neuron of Fig. 1E. Visually-evoked spikes were fewer on the long reaction time trials (top panel, in which the trials were sorted by saccadic reaction time), and this observation itself was likely one primary reason for why the visual response onset latencies were later on such trials (but see below for explicit analysis of this). Quantitatively, the Spearman correlation coefficient between trial-by-trial spike count in the visual burst epoch (Methods) and trial-by-trial saccadic reaction time for this example neuron was -0.6853 (p<0.0001) (Fig. 1E; bottom right panel). This suggests that weaker SC visual bursts were associated with later saccades, again as might be expected based on across-trial average measures ^15,17,19,20,41,42^. Importantly, this trial-by-trial correlation coefficient was quantitatively larger, in absolute value, than the correlation coefficient between trial-by-trial visual response onset latency and trial-by-trial saccadic reaction time in the same neuron (0.378). It was also quantitatively larger than the correlation coefficients seen in the two example V1 neurons of Fig. 1F, H (bottom right; -0.1313 and 0.0559 Spearman correlations between visual response strength and saccadic reaction time; p=0.3632 and 0.7286, respectively). These observations also held for the second example SC neuron of Fig. 1G (- 0.6402 Spearman correlation in this case; p<0.0001), as well as eight additional example SC and V1 neurons, shown in Fig. S2. Therefore, in the example neurons, visual response strength in the SC was the best predictor of saccadic reaction time variability when compared to SC visual response latency (and also to either visual response strength or latency in V1).

Across the population, these effects were robust (Fig. 2B). In the SC, the absolute values of the correlation coefficients in Fig. 2B were significantly larger than those associated with visual response onset latencies in Fig. 2A (U=480654, z=-6.14, p<0.0001, n_visual response latencies_=688, n_visual response strengths_=865, median_visual response latencies_=0.0804, median_visual response strengths_=-0.1543; Mann-Whitney U test comparing the absolute value of correlations for each correlation type – response onset latency versus response strength – across stimulus contrasts), and these effects also held at each contrast level individually after Bonferroni correction (except for 50% contrasts). Moreover, comparing the SC to V1 for visual response strength effects (Fig. 2B) revealed an essentially complete absence of correlations between trial-by-trial visual response strength and trial-by-trial saccadic reaction time in V1; in the SC, there were clear negative correlation coefficient values throughout all contrasts. We confirmed these observations statistically. Across all contrasts, V1 correlation coefficients were not significantly different from zero (W=663756, z=-1.14, p=0.2531, n=1656, Wilcoxon Signed-Rank test against zero). Moreover, at each contrast level individually, there were again no significant differences in the V1 correlation coefficients from zero after Bonferroni correction (10% contrast: W=25028, z=-1.93, p=0.0540, n=337; 20% contrast: W=39132, z=- 0.59, p=0.5569, n=402; 50% contrast: W=48175, z=-1.08, p=0.2781, n=452; 100% contrast: W=57195, z=1.04, p=0.2972, n=465). Finally, a Mann-Whitney U test comparing the SC to V1 correlation coefficients revealed stronger SC effects across all contrast levels (U=786061, z=- 17.59, p<0.0001, n_SC_=865, n_V1_=1658, median_SC_=-0.1543, median_V1_=-0.0026), and all individual comparisons within each contrast level were also significant after Bonferroni correction (p<0.0001).

The above results were also consistent at the individual monkey level (Fig. S3B, E, H). For the SC, each of the three tested monkeys showed clear negative population correlation coefficient values relating trial-by-trial visual response strength to trial-by-trial saccadic reaction time variability. And, in V1, even monkey F did not exhibit similarly strong negative population correlation coefficients as the SC values (Fig. S3E).

Finally, and for the sake of completeness, we additionally document in Fig. S4 the raw histograms underlying all of the data shown in this study so far, including an indication of the subset of the population for which significance was also reached at the individual-neuron level (Fig. 2). Consistent with the conclusions of Fig. 2, the SC histograms were the ones that deviated the most from zero, whether for visual response latency (Fig. S4A) or visual response strength (Fig. S4B).

Therefore, even though correlations to trial-by-trial saccade timing have been previously reported in the variability of V1 neurons (particularly in terms of visual response onset latency) ^23^, in our experiments, such correlations paled in comparison to those that we observed in the SC (Fig. 2A), and even more so when considering visual response strength (Fig. 2B). As we show next, this importance of SC visual response strength can be further appreciated when using statistical models combining both such strength and response latency in describing trial-by-trial saccadic reaction times.

### Superior colliculus visual response latencies interact with stimulus contrasts in accounting for behavioral variability

A relevant question given our results so far is whether visual response latencies in the SC have a real contribution to behavioral variability, or whether fewer spikes on slow trials (that is, weaker visual response strengths) simply mean later “first spikes” in the visual bursts (and subsequently later saccades). To answer this question, we fit a generalized linear mixed model (GLMM) for each stimulus polarity separately, predicting trial-by-trial saccadic reaction time as a function of SC visual response latency, SC visual response strength, and stimulus contrast, while neuron identity was included as a random intercept to account for across-neuron differences (Methods).

For dark contrasts, the model included a fixed intercept of 5.068 (SE=0.009), representing the expected log-transformed SRT when all predictors were at their mean. We only found significant main effects for visual response strength and stimulus contrast, but not for visual response latency (Wald Z-test; visual response latency: β=0.002, SE=0.001, z=1.388, p=0.165; visual response strength: β=-0.041, SE=0.002, z=-21.74, p<0.0001; stimulus contrast: β=-0.060, SE=0.001, z=-59.25, p<0.0001). However, the model also revealed a significant interaction between visual response latency and contrast (β=-0.006, SE=0.001, z=-6.94, p<0.0001), suggesting that for the weaker visual responses associated with low contrast stimuli, visual response latency in SC neurons still significantly explained some of the variance in saccadic reaction times, independently of visual response strength. This is consistent with the relatively higher correlation coefficient values seen in Fig. 2A for low versus high stimulus contrasts (in the SC part of the figure). In comparison, the interaction term between visual response strength and stimulus contrast did not significantly differ from zero (β=-0.0001, SE=0.001, z=-0.197, p=0.844).

We then performed subsequent single term deletion analyses, which removed each predictor along with its interactions. These analyses additionally revealed that even though there were small to moderate correlations present between the model predictors (visual response latency and visual response strength: r=0.328; visual response latency and stimulus contrast: r=0.473; visual response strength and stimulus contrast: r=0.122), the removal of each individual predictor and its associated interactions significantly worsened the model fit (Likelihood-ratio test; full model: AIC=285259, removing visual response latency: AIC=285309, χ^2^=54.58, df=2, p<0.0001; removing visual response strength: AIC=285724, χ^2^=469.61, df=2, p<0.0001; removing stimulus contrast: AIC=288618, χ^2^=3365.8, df=3, p<0.0001). This suggests that each predictor contributed unique information about trial-by-trial saccadic reaction time.

As for V1, a similar GLMM for dark contrasts revealed significant main effects of visual response latency and stimulus contrast, but not visual response strength (visual response latency: β=0.004, SE=0.001, z=4.897, p<0.0001; visual response strength: β=-0.001, SE=0.001, z=-0.633, p=0.527; stimulus contrast: β=-0.064, SE=0.001, z=-88.563, p<0.0001). As in the SC, only the visual response latency and contrast interaction was significant (β=- 0.005, SE=0.001, z=-7.564, p<0.0001), but not the visual response strength and contrast interaction (β=-0.001, SE=0.001, z=-0.860, p=0.390). The single term deletion analyses showed that in contrast to the SC, the removal of visual response strength as a predictor from the model did not significantly worsen it (Likelihood-ratio test; full model: AIC=403508, removing visual response latency: AIC=403597, χ^2^=92.803, df=2, p<0.0001; removing visual response strength: AIC=403506, χ^2^=1.56, df=2, p=0.459; removing stimulus contrast: AIC=411489, χ^2^=7986.6, df=3, p<0.0001).

Thus, SC, but not V1, visual responses, and especially their strengths, were the best predictors of the timing variability of foveating eye movements in our analyses. We next demonstrate how the differences observed so far between the SC and V1 were amplified even further when considering the visual responses of one specific functional class of visually-responsive SC neurons, and we later document the effects obtained with bright, rather than dark, luminance polarities.

### Greatest covariation with saccade timing in superior colliculus visual-motor neurons

The SC has two main functional classes of visually-responsive neurons: visual-only and visual-motor ^27,43,44^. The former appear like V1 neurons in the sense that they do not emit a saccade-related motor burst at the time of eye movement triggering (like in the example neuron of Fig. S2A), whereas the latter emit both visual and motor bursts (like in the example neurons of Figs. 1E, G, S2C, E, G). The visual neurons also receive the bulk of direct V1 inputs ^24–29^. We found that sensory-related correlations to behavioral variability in saccade times were always the highest in the visual responses of the SC’s visual-motor neurons, maximally differentiating these neurons from V1 neurons. This can be seen in Fig. 4A, B, in which we split the SC population of Fig. 2 according to the functional class of the encountered neurons (Methods). For both visual response onset latency (Fig. 4A) and visual response strength (Fig. 4B), visual-motor neurons clearly had the biggest effects. For example, at 50% and 100% (dark) contrasts, the visual response onset latency of visual SC neurons was barely correlated with trial-by-trial saccadic reaction time (50%: W=2937, z=1.22, p=0.2207, n=101; 100%: W=2656, z=1.40, p=0.1628, n=95; Wilcoxon Signed-Rank tests of the Spearman correlation coefficients against zero), but the correlation coefficients were several times larger for the visual-motor neurons (Fig. 4A; at each contrast level, correlation coefficients for visual-motor neurons were always significantly different from zero after Bonferroni correction, Wilcoxon Signed-Rank tests, p<0.0002). Across all contrasts, a Mann-Whitney U test revealed significantly larger correlation coefficient values for visual-motor than visual SC neurons (U=109302, z=-3.89, p=0.0001, n_visual_=348, n_visual-motor_=337, median_visual_=0.0572, median_visual-motor_=0.1024), with significant individual-contrast effects being also clear at 50% and 100% contrasts, after Bonferroni correction (again, as mentioned above, results for bright contrasts will be documented below).

**Figure 4.**
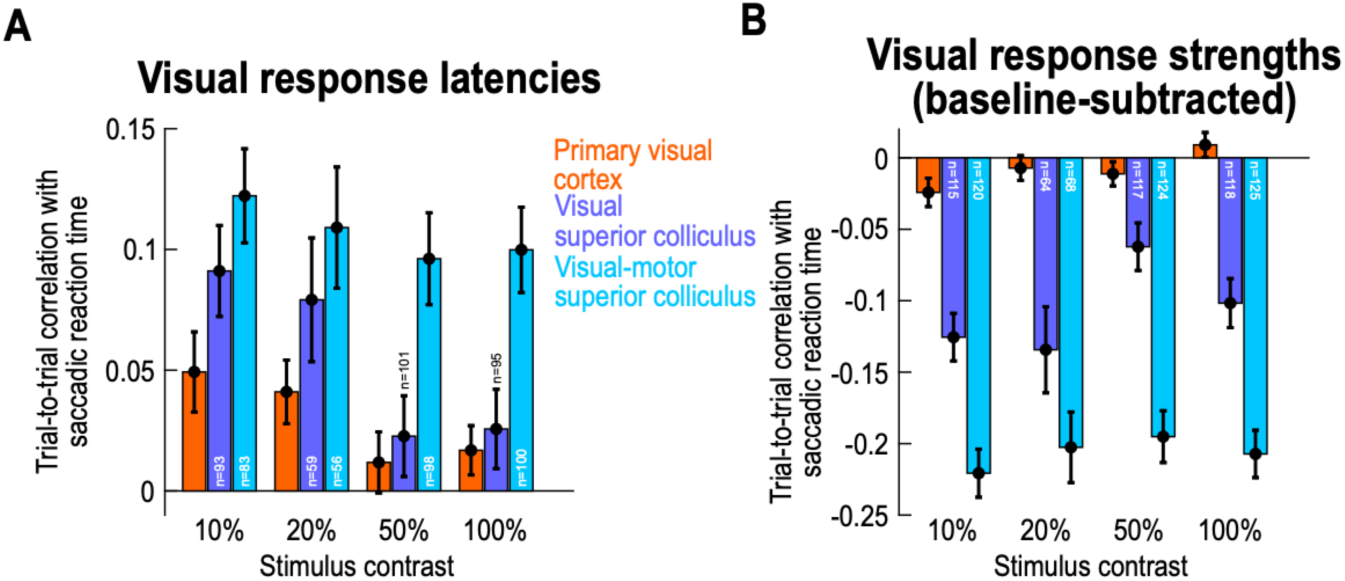
Strongest correlations with trial-by-trial eye movement timing in the visual responses of SC visual-motor neurons. **(A, B)** Same analyses as in Fig. 2 for the SC neuronal population (again for the dark contrasts), but now after separating the neurons according to whether they were visual-only or visual-motor (Methods). Visual-only neurons (purple) did not emit a saccade-related burst at saccade onset, whereas visual-motor neurons (cyan) emitted both a visual burst as well as a saccade-related one (as in the example neurons of Fig. 1E, G). For both visual response latencies (**A**) and visual response strengths (**B**), visual-motor neurons were better correlated with trial-by-trial saccade timing variability than visual neurons (that is, they had higher absolute values in their Spearman correlation coefficients). Moreover, the effect was much clearer for visual response strengths than visual response onset latencies. For example, correlations to saccade timing in the visual response strengths of visual-motor neurons were approximately twice as high as those of visual neurons (**B**). The orange bars show the V1 data (from Fig. 2) for comparison. Error bars denote SEM across neurons, and the numbers of neurons in each analysis are indicated in the figure (bars with missing numbers are replicates of other figures in which the numbers were given). Figure S5 shows individual monkey results, revealing a high level of consistency in the SC data across individuals, and Fig. S6 shows the raw histograms underlying all the SC data shown in the current figure.

Similarly, the correlations between trial-by-trial visual response strength and trial-by-trial saccadic reaction time were up to at least three times larger in visual-motor than visual SC neurons (Fig. 4B; U=204643, z=7.89, p<0.0001, n_visual_=414, n_visual-motor_=437, median_visual_=- 0.0958, median_visual-motor_=-0.2063; at each contrast individually, only 20% did not show a significant difference after Bonferroni correction). Thus, in terms of sensory processing, modulations in the visual responses of SC visual-motor neurons were the best predictors of subsequent behavioral variability ^19^.

Most critically, when comparing these results to those of V1 from Fig. 2, we confirmed how much stronger the difference between the SC and V1 became (the orange bars in Fig. 4 show the results from the V1 neurons for easier comparison). In fact, even though the SC’s visual neurons were the worst of the two functional SC neuronal types in the current analyses, they were still the better predictor of behavioral variability than V1 neurons (despite being the biggest recipients of direct V1 inputs ^25,27,29,45^). This conclusion can be visually inferred from comparing the purple to orange bars in Fig. 4A, B. Statistically, the correlation coefficients between visual response onset latencies and saccadic reaction times (Fig. 4A) were higher in the SC visual neurons than in V1 neurons across stimulus contrasts (U=262321, z=2.44, p=0.0149, n_SC_=348, n_V1_=1066, median_SC_=0.0572, median_V1_=0.0217; Mann-Whitney U test), and the correlation coefficients between visual response strengths and eye movement variability (Fig. 4B) were also higher in absolute value (that is, more negative) than in V1 (U=332183, z=8.90, p<0.0001, n_SC_=414, n_V1_=1658, median_SC_=-0.0958, median_V1_=-0.0026; Mann-Whitney U test). When testing at each individual contrast level, the correlation coefficient values were again stronger in SC visual neurons than in V1 neurons in terms of visual response strength (significant after Bonferroni correction, with p<0.003), but this was not the case for visual response onset latency. Thus, even though the SC’s visual neurons receive more of the direct V1 inputs than visual-motor neurons, they were still functionally distinct from V1 neurons. This is consistent with the idea that visual responses in the SC are reformatted versions of the inputs that they receive from the cortex and elsewhere ^24^.

The results of Fig. 4 for visual and visual-motor SC neurons were also all validated at the individual monkey level. This can be seen from Fig. S5: all three monkeys showed consistent results among each other, again validating the interpretation that, unlike in V1 (e.g. Fig. S3A, D), SC visual responses (and especially in the visual-motor neurons) were much more systematically related to behavioral variability than the other visual responses that we characterized in this study. In addition, and again for the sake of completeness, Fig. S6 shows all of the underlying raw histograms of the SC data of Fig. 4, supporting all of our conclusions from the population-level analyses.

### A phenomenon of putative anatomical depth in the superior colliculus’ layers

Our dichotomy of visual and visual-motor neurons above (Methods) suggests that, from the perspective of sensory drive, visual responses in the intermediate and deep anatomical SC layers are the ones that are most implicated in influencing the timing of foveating saccadic eye movements. This is because visual-motor neurons are consistently found in deeper SC layers than purely visual neurons ^46,47^. If so, then sorting all of our SC neurons according to their putative SC depths (independent of categorizing them individually into two discrete functional classes) should clarify the extent to which our SC effects were grounded in the underlying anatomy; such sorting should also confirm our interpretations from the results of Fig. 4 about visual-motor neurons.

To realize such sorting, especially since our animals are all still alive, we relied on a sensitive functional measure of SC neuron depth ^47–49^ (Methods). Specifically, using a delayed, visually-guided saccade task (Methods), we calculated, for each neuron, a visual-motor index (VMI) relating visual response strength (after stimulus onset) to saccade-related motor burst strength (at saccade onset). This index was +1 if there was no motor response above baseline but some visual response, and it was -1 if there was no visual response above baseline but some motor response; values in between +1 and -1 reflected the diversity expected from SC neuronal discharge across depths ^24,46–49^. We then binned all SC neurons as a function of their VMI values, and we plotted the average population correlation coefficient value (e.g. relating visual response strength to saccadic reaction time) in each functional “depth” bin.

Spearman correlation coefficient values between trial-by-trial SC visual response strengths and saccadic reaction times exhibited a step-like increase in absolute value for neurons with VMI’s less than zero (that is, for the deeper, more motor SC neurons). This result can be seen in Fig. 5 for the dark contrasts that we focused on so far in this study. The main panel in this figure shows that the population correlation coefficient values were generally negative across all VMI’s, consistent with Fig. 2 for visual response strength. However, the more motor neurons (negative VMI’s) had clearly stronger correlation coefficient values to behavior than the more visual neurons, and this effect was especially marked when VMI’s crossed from positive to negative values, a step-like change that is consistent with a layer-dependent effect. The insets in Fig. 5 show visual and motor bursts of three example neurons having different VMI values, to further illustrate our point. For the neuron with the most negative VMI (rightmost neuron in the figure), the visual response strength of the neuron was still large (reaching a peak of approximately 100 spikes/s, like the other two example neurons in the other two insets of the figure). The only difference is that the motor burst was now much stronger, making the visual burst appear small in the figure. This observation suggests that qualitatively similar visual response strengths in the SC (like across the three shown example neurons) can exhibit marked differences in their relation to behavioral variability: the deeper in the SC a visual response is encountered, the more correlated is this visual response to saccadic reaction time. Figure S7 shows that similar conclusions could also be reached when considering visual response latency in the SC instead of visual response strength, but the effects were expectedly weaker (given all of our results observed so far, and given the noisier nature of detecting visual response latencies of individual neurons). The figure also shows the visual response strength results with bright contrasts (Fig. S7C), again showing a step-like discontinuity as we traversed the putative depths of the SC, just like in Fig. 5.

**Figure 5.**
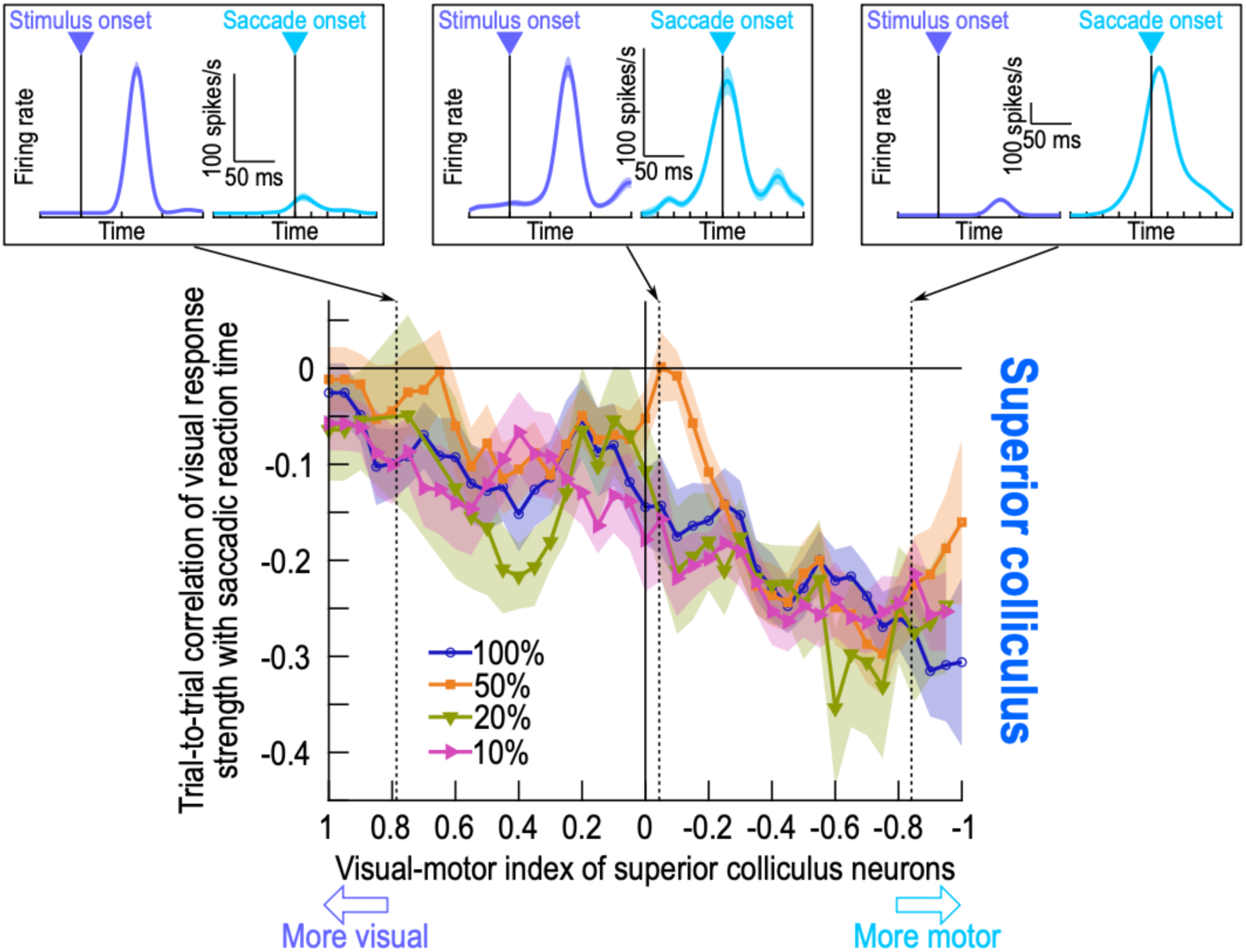
Dependence of the SC results on the functionally-defined depths from which the neurons were recorded. The y-axis of the main panel shows the Spearman correlation coefficient values when relating trial-by-trial SC visual response strength to trial-by-trial saccadic reaction time; the x-axis shows the visual-motor indices (VMI’s) of the recorded SC neurons. Indices on the x-axis closer to 1 (left side of the axis) indicate more visual neurons, and indices closer to -1 (right side of the x-axis) indicate more motor neurons. For each contrast level (for the dark stimuli in this figure), there was a step-like change in correlation coefficient values, with the visual responses in the more motor neurons (deeper SC layers) showing stronger negative correlations to saccadic reaction time. Thus, even without having to functionally classify neurons as in Fig. 4, we confirmed that visual-motor SC neurons had the strongest effects when linking visual response strength to behavior. Note that the insets above the main panel show example visual and motor responses (Methods) from three SC neurons occupying different regions of the VMI space. The leftmost neuron was predominantly visual (VMI approximately 0.8), whereas the rightmost neuron was predominantly motor (VMI approximately -0.8). Note that the visual response strength of this more motor neuron was still peaking at around 100 spikes/s, which is similar to the visual response strengths of the two other shown neurons. Thus, qualitatively similar visual response strengths can show quantitatively different correlation coefficient values to behavior, simply as a function of their putative depth in the SC. Error bars denote SEM across neurons in a given VMI bin. And, Fig. S7 shows similar analyses for the visual response latencies of our SC neurons, and also for the results (both on response latency and response strength) obtained with bright visual stimuli.

Therefore, at least from the perspective of sensory-driven modulations, visual responses in intermediate and deeper SC layers (as functionally identified using VMI analyses) carry an especially relevant functional meaning for explaining trial-by-trial variability in the timing of foveating saccadic eye movements.

### Superior colliculus pre-stimulus activity is lower for later saccades

The activity of SC neurons can additionally reflect a readiness to move the eyes even before stimulus onset ^42,50,51^. Therefore, and for completeness, we next measured pre-stimulus (baseline) activity and checked how well it predicted trial-by-trial variability in saccade timing. For V1, there were no consistent effects (Fig. 6A), in line with all of our results above. Specifically, a Wilcoxon Signed-Rank test revealed no difference from zero across contrasts (W=366580, z=-0.47, p=0.6360, n=1220), although for only the 20% contrast individually, there was a significant positive correlation after Bonferroni correction (W=26332, z=3.42, p=0.0006, n=292). On the other hand, and in agreement with the prior SC literature on motor readiness ^42,51^, SC pre-stimulus activity was systematically weaker for slower saccadic reaction times (Fig. 6A; W=92325, z=-9.26, p<0.0001, n=774; Wilcoxon Signed-Rank test against zero across contrasts; comparisons to zero at each contrast level individually were also significant after Bonferroni correction, with p<0.0001). Moreover, the difference between the SC and V1 effects was statistically significant (U=949601, z=-8.21, p<0.0001, n_SC_=865, n_V1_=1658, median_SC_=-0.0407, median_V1_=0; Mann-Whitney U test comparing SC to V1 across contrasts; comparisons across the brain areas for each contrast level individually were also significant after Bonferroni correction, with p<0.0013). Therefore, we identified another substantial difference between the SC and V1, this time with respect to pre-stimulus state.

**Figure 6.**
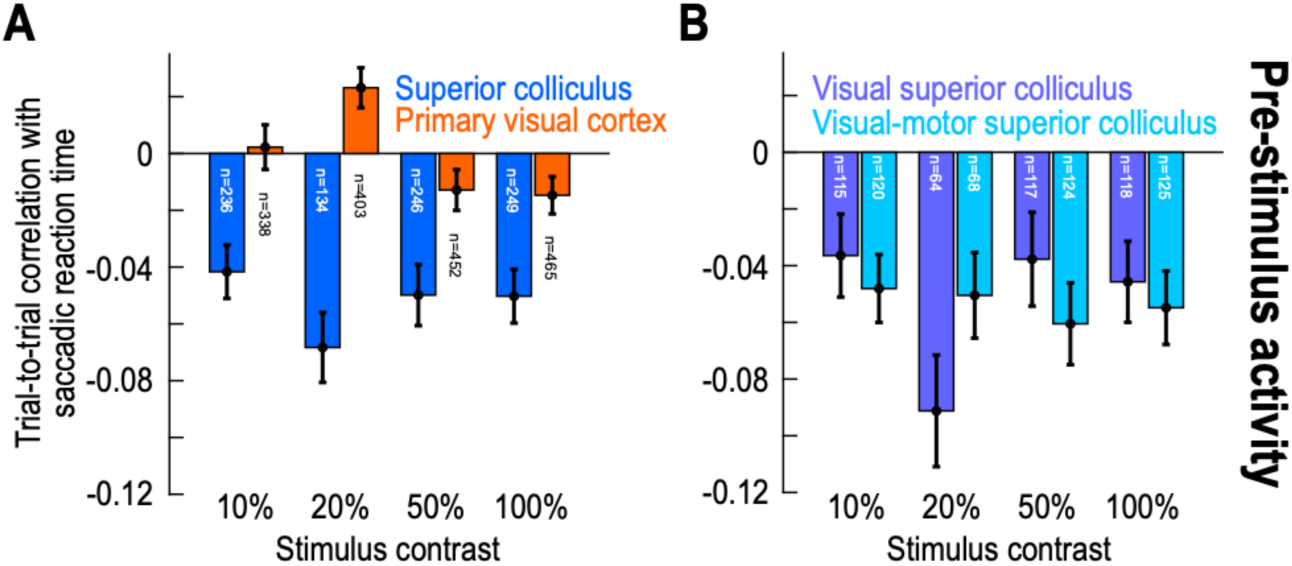
An additional correlation with saccade timing in the SC’s, but not V1’s, pre-stimulus (baseline) neuronal activity. **(A)** Similar to the analyses in Fig. 2, again for the dark contrasts. However, here, we measured the correlation coefficients between trial-by-trial pre-stimulus (baseline) activity (Methods) and trial-by-trial saccade timing. In the SC, there was a clear negative correlation at all tested contrasts: higher pre-stimulus activity was associated with faster saccadic reaction times. This was not the case in V1. **(B)** This effect was similar regardless of whether the SC neurons were visual-only or visual-motor. Thus, in addition to visual response latency and strength in the SC (Figs. 1–5), the pre-stimulus state of the SC can much more strongly influence saccadic reaction times than the V1 pre-stimulus state. Error bars denote SEM, and the numbers of neurons in each analysis are indicated in the figure.

We likewise repeated the same analysis of Fig. 6A, but now separately for either visual or visual-motor SC neurons. There were similar relationships between pre-stimulus SC state and saccadic reaction time for both types of neurons (Fig. 6B). Specifically, there was no effect of neuron type (U=178007, z=0.46, p=0.6466, n_visual_=414, n_visual-motor_=437, median_visual_=-0.0435, median_visual-motor_=-0.0343; Mann-Whitney U test across contrasts; comparisons at each contrast level individually also revealed no significant differences after Bonferroni correction). Thus, pre-stimulus SC state was, in general, an important determinant of saccade timing variability.

Taken together, our results so far reveal a striking difference in the relationship between SC and V1 stimulus-related neuronal activity to trial-by-trial variability in saccadic reaction times, with our take-home message being: there is a much higher covariation with foveation timing by SC than V1 stimulus-aligned activity. Figure 7 succinctly summarizes this message. Here, we normalized each neuron’s firing rate curve before averaging across neurons (Methods), and we compared the fastest and slowest third of all trials (in terms of saccadic reaction time) within a given session. In the SC (Fig. 7A), faster saccadic reaction time trials were associated with: (1) higher pre-stimulus (baseline) activity; (2) earlier visual response onset latency; (3) and higher visual response strength. Moreover, note that the visual response strength effect was not simply inherited from the pre-stimulus (baseline) effect; in fact, in all of our visual response strength analyses above and in the remainder of this study, we always measured the baseline-subtracted visual response strength (Methods) in order to exclude the impact of pre-stimulus activity. As for V1, neuronal activity did not differ on the fast and slow trials at all (Fig. 7B), consistent with the results of Figs. 1–6, but inconsistent with previous results on visual response onset latency ^23^. In what follows, we describe our investigations of whether and how stimulus luminance polarity (bright versus dark) or V1 neuronal classes might have qualified (at least ever so slightly) the above conclusions.

**Figure 7.**
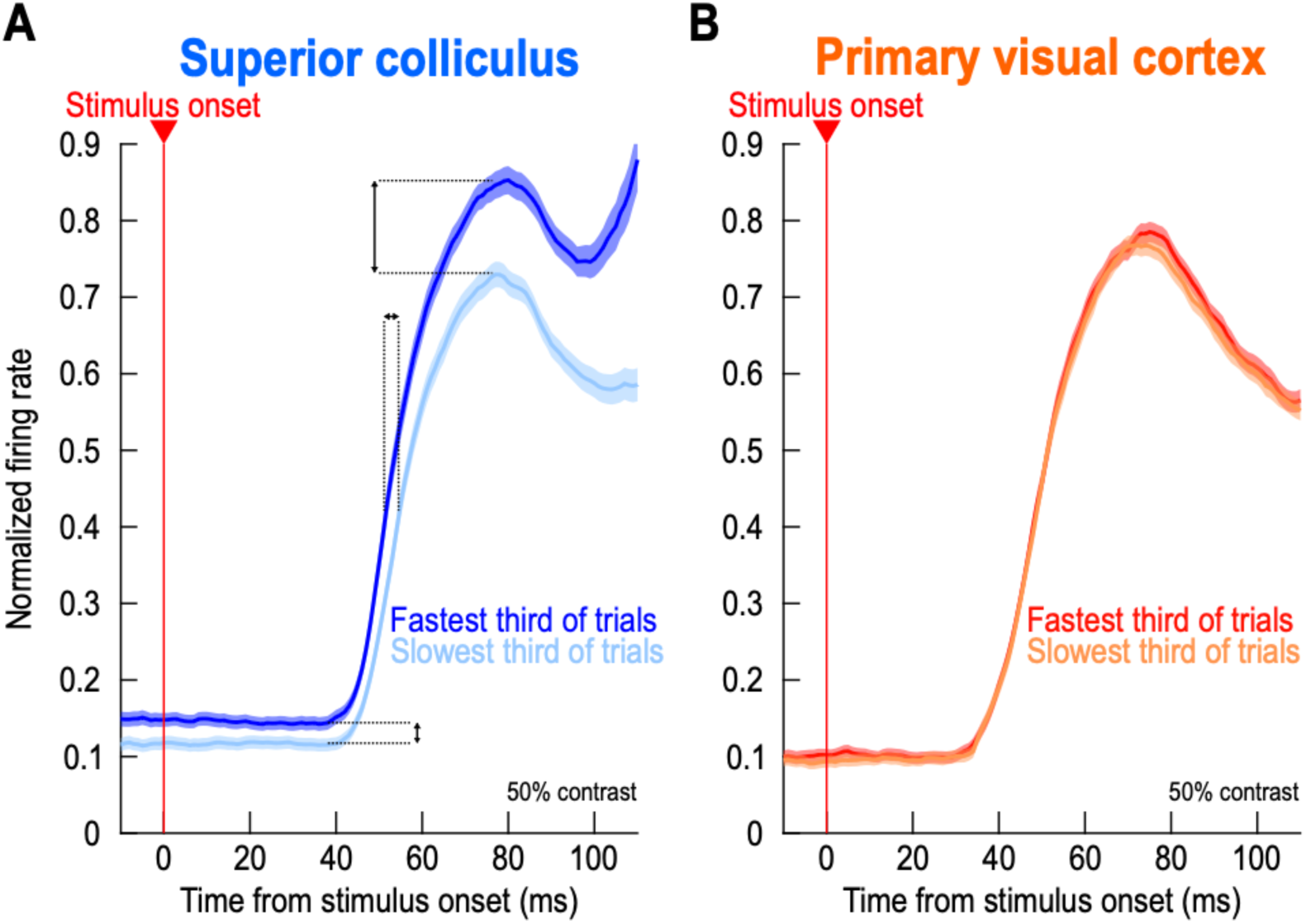
Summary of the differences between the SC and V1 with respect to the timing of foveating saccadic eye movements. **(A)** For an example stimulus level (50% dark contrast), we normalized each neuron’s trial-by-trial activity to the peak of its average visual response across trials (Methods). We then measured neuronal activity on either the fastest or slowest third of all trials for the neuron. Across neurons in our entire SC population, trials with the fastest saccadic reaction times were consistently the ones with elevated pre-stimulus (baseline activity), early visual latencies, and strong visual burst amplitudes. **(B)** On the other hand, V1 activity did not predict saccadic reaction times. Error bars denote SEM across neurons, and the numbers of neurons are the same as those that went into the visual response strength analyses of Fig. 2.

### Similar results for bright stimuli

Besides presenting negative luminance polarity stimuli (Figs. 1–7), we also included similar Weber contrast levels, but with stimuli that were now brighter than the surrounding background (Fig. 1D; Methods) ^18^.

For the SC, there were no differences in correlation coefficient values between visual response onset latency and trial-by-trial variability in saccadic reaction times (Fig. 8A; U=466079, z=0.38, p=0.7038, n_brights_=658, n_darks_=688, median_brights_=0.0821, median_darks_=0.0804; Mann-Whitney U test across contrasts; comparisons within each contrast level individually were also not significant). In terms of visual response strengths (Fig. 8B), there likewise were no significant differences between dark and bright contrasts (U=724830, z=-1.85, p=0.0638, n_brights_=854, n_darks_=865, median_brights_=-0.1347, median_darks_=-0.1543; Mann-Whitney U test across contrast levels), and none of the comparisons within each contrast level were individually significant after Bonferroni correction.

**Figure 8.**
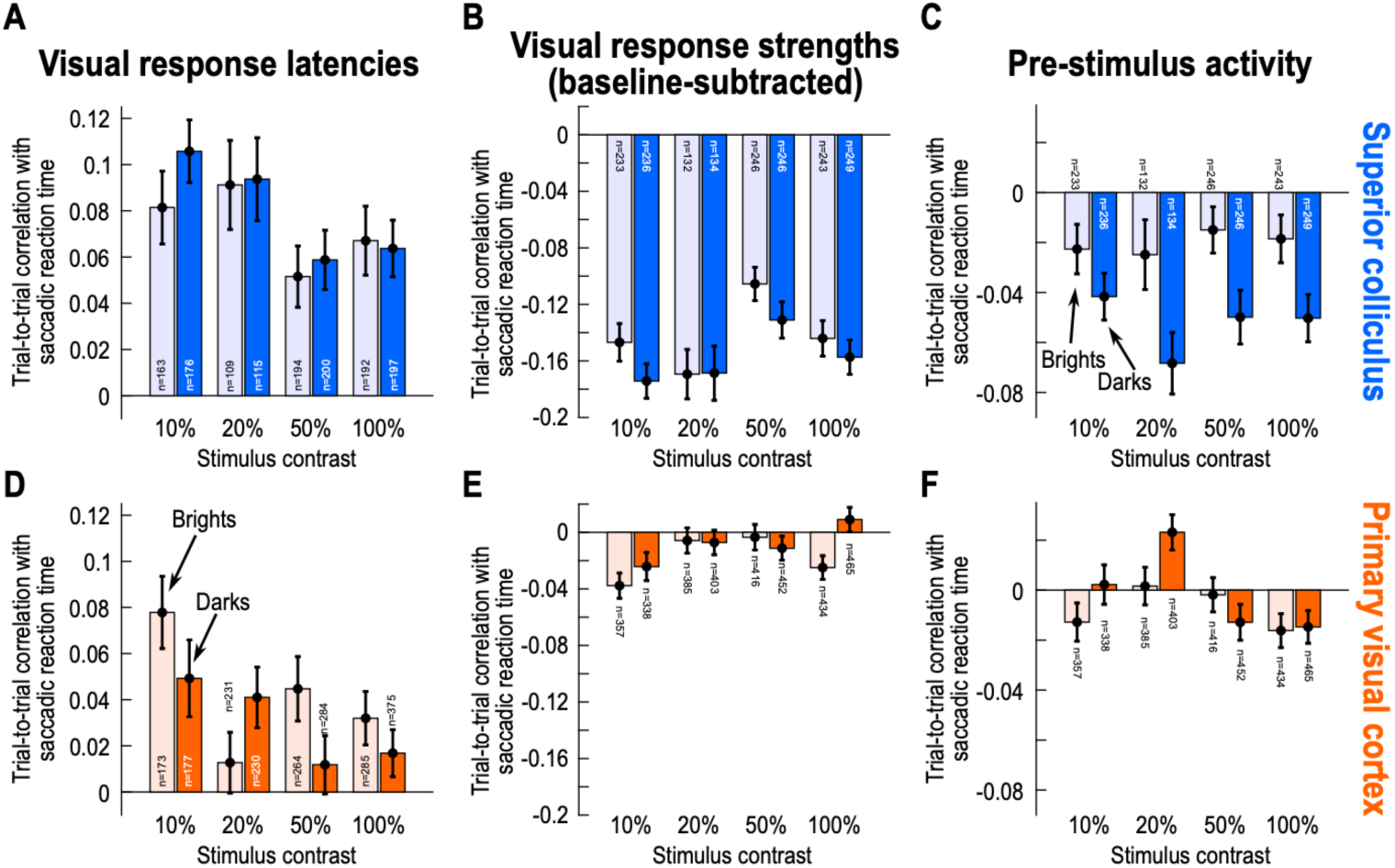
Mildly weaker effects for bright contrasts in the SC, but mildly stronger visual response latency correlations with saccade timing for bright contrasts in V1. **(A-C)** Same analyses as in Figs. 2, 6A for the SC neurons in our population. The saturated colors show correlations with trial-by-trial saccade timing (of the relevant neuronal parameter in each panel) for the dark contrast stimuli in our experiments (same as in Figs. 2, 6A). The faint colors show results for bright contrasts (stimulus onsets brighter than the surrounding gray background; Methods). There were generally weaker correlations for the bright contrasts, especially in pre-stimulus (baseline) activity (**C**). **(D-F)** Similar analyses for V1. In this case, only for visual response timing (**D**) was there a trend for higher correlation with trial-by-trial saccadic reaction time for bright than dark contrasts. Error bars denote SEM, and the numbers of neurons in each analysis are shown in the figure.

In contrast, pre-stimulus SC state did substantially differ between bright and dark stimuli (Fig. 8C). For bright stimuli, variability in pre-stimulus SC state was less related to variability in trial-by-trial saccadic reaction times than for dark stimuli, and this effect was significant (U=701280, z=-4.14, p<0.0001, n_brights_=854, n_darks_=865, median_brights_=-0.0037, median_darks_=-0.0407; Mann-Whitney U test comparing brights and darks across contrasts). At the individual contrast levels, the statistical comparisons between bright and dark stimuli were significant after Bonferroni correction at 20% and 50% contrasts (p<0.011). These observations add to evidence that the recruitment of SC visual bursts can be different between bright and dark stimuli ^18,52^ (also see Fig. 9 below for further evidence). Indeed, when we measured the correlation coefficient values between trial-by-trial pre-stimulus state and the trial-by-trial visual response onset latency of SC neurons, there was no difference between dark and bright stimuli (Fig. S8A; U=420300, z=0.16, p=0.8729, n_brights_=633, n_darks_=652, median_brights_=-0.1168, median_darks_=-0.1060; Mann-Whitney U test comparing brights and darks across contrasts). That is, for both types of stimuli, weaker pre-stimulus SC activity was associated with later visual response onset latencies, and by a similar amount whether the appearing stimuli were bright or dark. Thus, the negative correlations to behavioral variability in Fig. 8C were not directly explained by variability in visual response onset latency in the SC, and likely reflected intrinsically different mechanisms for generating visual responses to bright stimuli ^18^.

**Figure 9.**
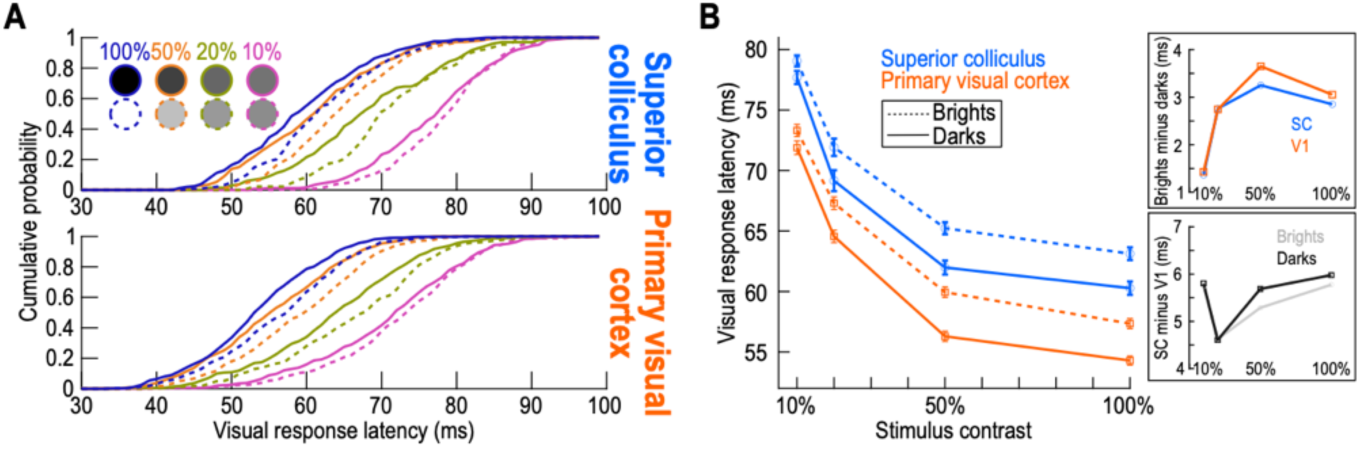
Visual responses are generally later for bright than dark contrasts in both the SC and V1. **(A)** Cumulative distributions of the average visual response latency (across trials) of each neuron in our SC (top) and V1 (bottom) populations. Low contrasts were associated with generally longer average visual response latencies in both areas than high contrasts, as expected. Additionally, at each contrast level, dark stimuli had earlier visual response latencies. **(B)** Summary statistics of the results in **A**. V1 visual responses (orange squares) were earlier than SC (blue circles) visual responses (by 5-6 ms; see bottom right inset); moreover, in both areas, bright contrasts had later visual responses than dark contrasts (by 1-3 ms; see top right inset). The contrast-dependence of bright and dark latency differences in the SC (top right inset) quantitatively and qualitatively replicates our earlier results (primarily from a gaze fixation task) ^18^. Error bars in the main panel of **B** denote SEM across neurons, and the numbers of neurons included at each contrast level are the same as those for the visual response strength analyses of Fig. 2.

In terms of SC neuron type, the pre-stimulus effect seen in Fig. 8C was slightly clearer in visual than visual-motor SC neurons (Fig. S9). Thus, just like in Figs. 4, 5, there were still potential functional differences between SC visual and visual-motor neurons when exploring effects of stimulus luminance polarity.

In V1, for visual response onset latency, there was no statistically significant difference in the Spearman correlation coefficient values relating to behavioral variability between bright and dark stimuli (Fig. 8D; U=1056948, z=-1.51, p=0.1317, n_brights_=953, n_darks_=1066, median_brights_=0.0361, median_darks_=0.0217; Mann-Whitney U test across contrasts). This was also true in the individual pairwise comparisons between brights and darks at each contrast level (p>0.08 across all individual tests).

In terms of visual response strengths in V1 (Fig. 8E), there were larger absolute values of correlations to behavioral variability for brights than darks (U=2751036, z=2.09, p=0.0364, n_brights_=1592, n_darks_=1658, median_brights_=-0.0173, median_darks_=-0.0026; Mann-Whitney U test across contrasts), but for the individual contrast levels, only 100% contrasts revealed a difference between bright and dark stimuli after Bonferroni correction (p=0.0067; Mann-Whitney U test). This suggests only moderately clearer, if any, covariation with behavioral variability in V1 neurons when the visual stimuli involved were brighter than the background.

And, for pre-stimulus (baseline) activity (Figs. 8F, S8B), there were no differences in the correlation coefficient values between bright and dark stimuli in V1 (U=2728841, z=1.2744, p=0.2025, n_brights_=1592, n_darks_=1658, median_brights_=0, median_darks_=0).

Finally, we also performed all of our GLMM analyses that we completed above, but this time one more time for the bright stimuli. In the SC, rebuilding the GLMM for bright contrasts showed that in addition to visual response strength and stimulus contrast, visual response latency was a statistically significant main effect in this model (Wald Z-test; visual response latency: β=0.008, SE=0.001, z=7.909, p<0.0001; visual response strength: β=-0.027, SE=0.002, z=-16.61, p<0.0001; stimulus contrast: β=-0.056, SE=0.001, z=-66.310, p<0.0001). Furthermore, not only the visual response latency and contrast interaction term significantly contributed to the model (β=-0.009, SE=0.001, z=-10.053, p<0.0001), but the visual response strength and contrast interaction term did as well (β=-0.002, SE=0.001, z=-2.147, p=0.0318). The single term deletion analyses showed that each predictor contributed unique information about saccadic reaction time (Likelihood-ratio test; full model: AIC=274018, removing visual response latency: AIC=274197, χ^2^=182.96, df=2, p<0.0001; removing visual response strength: AIC=274247, χ^2^=232.96, df=2, p<0.0001; removing stimulus contrast: AIC=278020, χ^2^=4007.3, df=3, p<0.0001).

For the bright contrasts in V1, like for the dark contrasts, we found significant main effects of visual response latency and stimulus contrast, but not visual response strength (visual response latency: β=0.006, SE=0.001, z=7.181, p<0.0001; visual response strength: β=-0.002, SE=0.001, z=-1.832, p=0.067; stimulus contrast: β=-0.047, SE=0.001, z=-66.749, p<0.0001). As in the SC for bright contrasts, both the visual response latency and contrast interaction (β=-0.006, SE=0.001, z=-8.738, p<0.0001) and the visual response strength and contrast interaction (β=-0.003, SE=0.001, z=-3.891, p<0.0001) were significant. The single term deletion analyses showed that the removal of any of the predictors from the model significantly worsened it (Likelihood-ratio test; full model: AIC=379805, removing visual response latency: AIC=379951, χ^2^=149.76, df=2, p<0.0001; removing visual response strength: AIC=379821, χ^2^=19.905, df=2, p<0.0001; removing stimulus contrast: AIC=384604, χ^2^=4804.9, df=3, p<0.0001).

### Average visual response onset latencies are similarly affected by brights and darks in the two brain areas

As alluded to above, the recruitment of visual responses, in both the SC ^18,52^ and V1 ^53–55^, can be fundamentally different between bright and dark stimuli. Consistent with this, across-trial average visual response onset latencies in our current data were always longer for bright than dark stimuli, and at all tested contrasts (Fig. 9). Even though our primary focus was on trial-by-trial variability, these analyses are important to document here because they compare, for the first time, the visual response latencies from both brain areas in the very same animals and with the very same stimuli. Interestingly, the contrast-dependent difference between bright and dark average visual response onset latencies (Fig. 9B; top right panel) was similar between the SC and V1, and the SC data quantitatively replicated our previous observations ^18^. Additionally, Fig. 9 shows that SC visual responses were always 5-6 ms later than V1 visual responses, on average (Fig. 9B; bottom right panel).

### Lack of difference across cortical layers and putative cortical cell types

Because of the striking difference in how SC and V1 visual neuronal activity variability related to behavior (Figs. 1–8), we next wondered whether individual subsets of V1 neurons might have shown substantially larger effects than in our summary analyses.

We first considered cortical depth. Using current source density (CSD) analysis ^56^, we estimated the input layers of V1 from each session, and we then classified our V1 neurons as being either in the superficial or deep cortical layers (Methods; Fig. S10A, B). For both visual response onset latencies and visual response strengths, the V1 correlations to behavioral variability were similar across cortical depths (Fig. S10C-F). We did not perform similar analyses for V1 pre-stimulus state, since V1 baseline activity was anyway already much weaker than SC pre-stimulus activity (Fig. S11A), and also because all of our other analyses did not show effects with V1 pre-stimulus state. Moreover, pre-stimulus (baseline) V1 activity was similar for the deep and superficial cortical layers (Fig. S11B).

Finally, for a subset of our V1 neurons, we additionally had suitable spike waveforms, allowing us to classify the neurons by their waveform widths (Methods). We used a similar approach to earlier studies, which classified putative excitatory and inhibitory early visual cortical neurons based on their spike waveform widths ^57^ (Methods). Once again, there were no differences between putative excitatory (broad-spiking) and putative inhibitory (narrow-spiking) V1 neurons (Fig. S12).

Thus, the large difference that we observed between the SC and V1 in our data set (e.g. Fig. 7) was a general property of the two brain areas, and did not depend on specific cortical subunits.

## Discussion

We were motivated by the idea that multiple brain areas can exhibit qualitatively similar short-latency visual responses to appearing stimuli, raising the question of why such apparent parallelism is necessary at all. This question is even more pressing in the case of the SC and V1, especially because the SC receives a direct feedforward projection from V1 ^24–30^, and also because previous work ^15,23^ has shown that both areas might be relevant for saccade timing variability. By studying both areas within the same animals and with the same visual stimuli, we could identify important functional differences between their visual responses.

We found that visual response strength in the SC was a better predictor of trial-by-trial saccadic reaction time than visual response latency in the same neurons, and we also found that both visual response strength and visual response latency in the SC were the better predictors of behavioral variability when compared to the same neuronal response parameters obtained from V1 neurons. This suggests that SC visual responses were not simply inherited versions of their V1 counterparts. In addition, our GLMM’s revealed that SC visual response latency could still contribute to the behavioral variability, independently of visual response strength, especially at low stimulus contrasts. Finally, all of these effects were amplified when we analyzed the visual responses of visual-motor SC neurons, residing in the deeper layers of the SC, further supporting the view that these neurons are critical for sensory-motor transformations in eye movement behaviors.

In terms of absolute latencies, we found that neurons in V1 “detect” new visual stimuli 5-6 ms earlier than SC neurons (Fig. 9). However, SC neurons much more directly, and much more effectively, drive behavior (Figs. 1–8). This is consistent with the proximity of the SC to the motor output ^58^, and it is also consistent with the role of the SC in salience representation ^59–62^. Thus, V1 senses the environment, but the SC evaluates and acts on it.

It is interesting to contemplate here about how our results relate to neuronal variability in general, both cortically and sub-cortically. Specifically, because saccadic reaction time variability is generally high (Figs. 1, S13), does the higher correlation that we observed in the SC to such variability automatically imply that SC visual responses are intrinsically much less reliable than V1 visual responses? In our data, this was decidedly not the case. For example, for each neuron in either V1 or the SC, we measured the standard deviation across trials of visual response onset latency (Fig. 10A, B) or visual response strength (Fig. 10C, D) within any given session. Across the population, and perhaps unexpectedly, we found systematically higher variability across trials in V1 than in the SC. Thus, SC visual responses were actually more reliable than V1 visual responses. This is somewhat surprising given prior suggestions that V1 visual responses are highly reliable ^63^, and it suggests that there are other sources of variability in V1 beyond those that ultimately get reflected in the final eye movement behavioral variability. These other sources are unlikely to include changes in stimulus position in the V1 RF’s from trial to trial, because we used a relatively big stimulus for V1 RF sizes, and also because we did not see much evidence of strong modulations due to, say, microsaccades in V1 (also see below). It would be interesting in the future to understand the functional advantages of such higher response reliability that we observed in the SC in Fig. 10.

**Figure 10.**
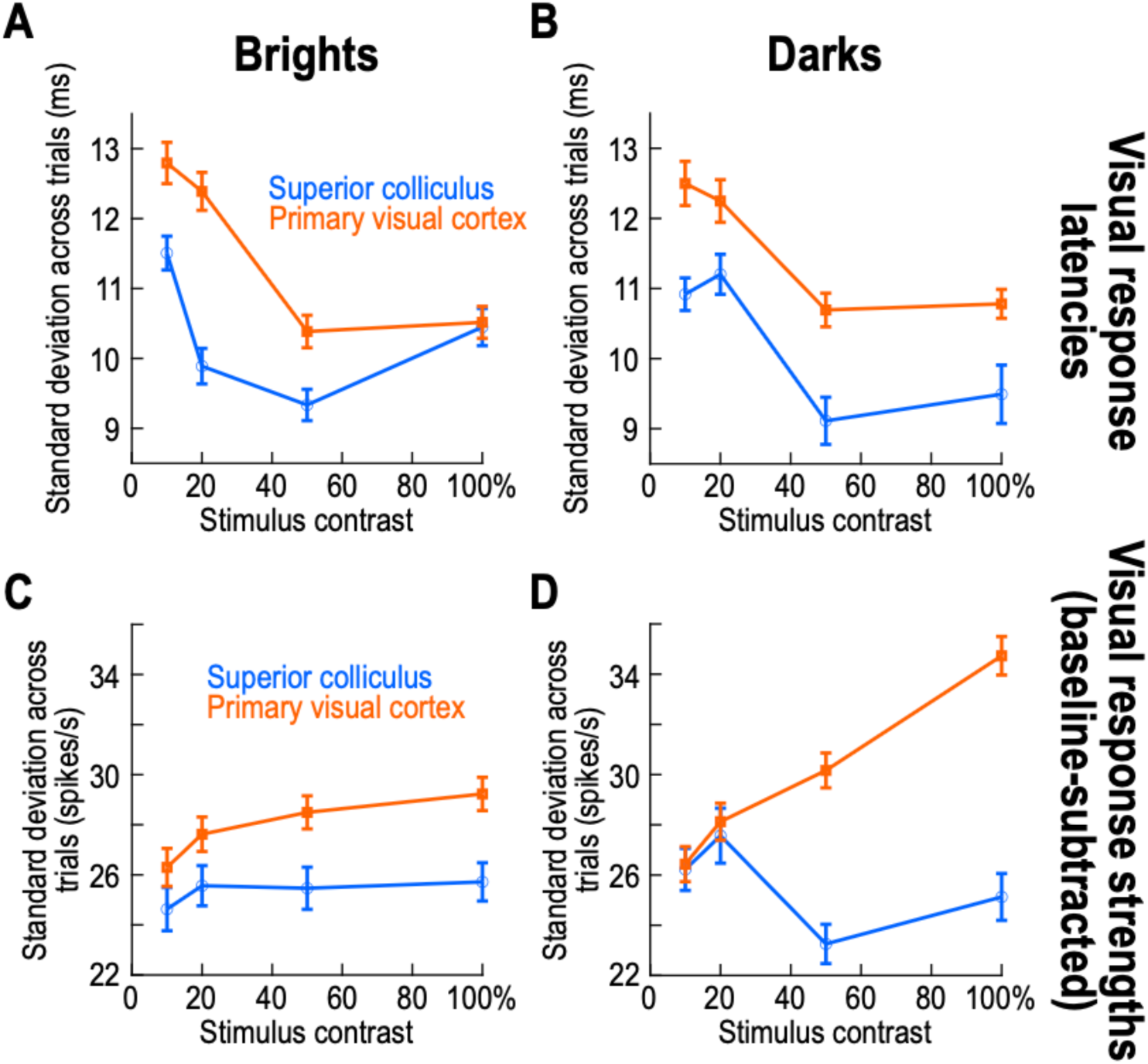
Higher response reliability in SC than V1 visual responses, across contrasts and luminance polarities. **(A, B)** For each neuron in either the SC (blue circles) or V1 (orange squares), we measured the standard deviation of visual response onset latencies across trials within a given session. Then, we summarized this measure across all neurons in the population, by plotting the mean and SEM ranges. For both positive (**A**) and negative (**B**) luminance polarity stimuli, SC visual responses were more reliable than V1 visual responses across trial repetitions. Thus, given that V1 correlations to behavioral variability were low, and given that behavioral variability was itself very high (Fig. S13), V1 visual responses have other sources of variability in them, which are not ultimately reflected in the behavioral eye movement output variability. **(C, D)** Same as **A**, **B** but for baseline-subtracted visual response strength variability. Dark stimuli had a monotonic increase in V1 variability with increasing contrast (in terms of visual response strength), which suggests that contrast sensitivity in V1 is different between bright and dark stimuli. Error bars indicate SEM across neurons.

Our results also allow making concrete predictions about further differences between SC and V1 visual responses. For example, a well-established reason for variability in SC visual responses, and subsequent saccadic behavior, is the occurrence of peri-stimulus microsaccades. Such microsaccades significantly alter SC visual sensitivity ^19,20,52,64^, and with direct consequences on behavioral timing variability ^19,20,65^. The fact that we observed weaker V1 than SC trial-by-trial covariation with subsequent behavioral timing suggests that peri-stimulus microsaccades might have weaker influences on V1 than on the SC. This is a fundamentally important prediction because it can allow identifying the brain loci for robust perceptual and cognitive phenomena, such as saccadic suppression of perceptual sensitivity ^66,67^ or covert visual attentional performance ^68,69^, which are known to be strongly influenced by microsaccades. In fact, the modest positive correlations to behavioral variability that we observed in V1 for bright stimuli (e.g. Fig. 8) can even allow predicting a dichotomous influence of saccadic suppression (due to peri-stimulus microsaccades) on either bright or dark visual stimuli in V1; it could be the case that dark contrasts are immune to saccadic suppression in V1 but not in the SC, consistent with existing preliminary evidence ^70^. This would be very interesting to further investigate, especially because in other species, there seems to indeed be a dependence of saccadic suppression on stimulus luminance polarity and behavioral context ^71,72^.

It would also be interesting to analyze simultaneously recorded neurons in a paradigm like ours, to see whether populations of neurons covary during the behavior. For example, if we were to sort trials by saccadic reaction times and analyze the visual response strengths of two simultaneously recorded SC neurons, would the two neurons exhibit similar rank ordering of their visual response strengths as a function of saccadic reaction time, or would other sources of variability dominate the pairwise correlations of the neurons’ activity? Similarly, one could ask these same questions for pairs of V1 and SC neurons, and even for saccade landing accuracy and precision as opposed to saccade timing.

Concerning the large differences that we saw between SC and V1 trial-by-trial correlations to behavior (in terms of sensory-related activity), our observations confirm assertions that visual responses in the SC are not merely a copy of V1 visual responses, but are instead reformatted versions that are better suited for the SC’s functional roles ^24^. In other words, assuming that the SC and V1 have distinct functional roles in perception, cognition, and behavior, which is a fair assumption, then their individual visual response properties should support their respective individual roles. Indeed, SC neurons prefer low spatial frequencies ^17^, whereas V1 neurons are more band-pass at similar retinotopic eccentricities ^73,74^, and this is directly relevant for saccade timing ^17,19^. Similarly, SC neurons prefer (in visual sensitivity, resolution, and potential over-representation) the upper visual field ^75,76^, whereas V1 might even over-represent at least parts of the lower visual field, such as the lower vertical meridian ^77–79^, and this is again relevant for the use of eye movements to achieve remote sensing ^75,80^. Finally, SC neurons do not directly copy the preference of V1 neurons for dark contrasts ^18^, and exhibit a dissociation between response strength and response timing that likely matters for how quickly the SC can trigger eye movements ^17,18^. Our present results add that response variability in both brain areas is also very different, but in a perfectly meaningful way when it comes to how visual responses in each of the areas may be subsequently used by downstream neurons.

Of course, it would be interesting to additionally investigate monkeys like our monkey F in Fig. S3D. This monkey seemed to be more reflexive than the other two monkeys (Fig. S3F), and it also showed positive correlation coefficient values between V1 visual response latencies and saccadic reaction times. This observation, coupled with the mildly stronger correlations that we saw in V1 for bright stimuli (Fig. 8), can be a way to reconcile our results with those of the previous V1 investigation of the same topic ^23^. There, the authors used Gabor gratings as the saccade targets, which had both positive and negative polarity stimulus subcomponents. Therefore, it could be that their results were dominated by the positive luminance polarity components of the gratings. Having said that, we cannot directly quantitatively compare our V1 results to their data due to the different stimuli used. Moreover, in that earlier study, the saccade target could appear at one of two or four possible stimulus locations on the display across trials ^23^, whereas we had one location (but different stimulus timings and appearances).

Finally, just like we cannot directly compare our V1 results to those of ref. ^23^, we also cannot quantitatively compare our correlation coefficient values in the SC with previous SC measures of correlations between visual responses and saccade timing ^15,16^. This is because we used an immediate, visually-guided saccade task in our experiments, whereas this prior work used either a delayed or gap saccade task. In the delayed saccade task, the animal sees the eccentric target for a long time and has to withhold orienting towards it until instructed to do so, and in the gap saccade task, the initial fixation target is extinguished before the appearance of the eccentric target; in both cases, there are well-documented increases in SC motor preparation that are not normally observed in the immediate saccade task. Moreover, in our case, even though the expected target location was known to the animals, there was substantial uncertainty about both the timing and the visibility of the upcoming visual stimulus. Thus, a direct quantitative comparison is decidedly not possible, and that is why we elected to compare our two brain areas directly with the very same task in the same animals. Interestingly, and on a related note, it might additionally be wondered what the removal of the foveated fixation target (together with the eccentric target appearance) does to behavioral variability in our task. This is an ongoing question, but our hypothesis is that foveation timing variability is much more robustly related to visual responses in the SC, than to the effects of foveal target removal at the initial fixation position.

### Limitations of the study

In this study, the target location was fixed throughout each session, which allowed us to measure neuronal responses on every collected trial. However, in the future, it would be important to also have an unpredictable target location across trials, so that the saccades are truly reflexive. This means that we might not be able to collect neuronal responses to target onset on every single trial, but it would help eliminate target location predictability as a variable for potentially explaining part of our results.

It would also be worthwhile in the future to consider images of real-life objects as the saccade targets ^81,82^, and to even record SC and V1 activity during free viewing of natural scenes. This can help clarify the ecological relevance of our SC observations.

Finally, the nature of the computational transformation happening from V1 to SC visual responses is unknown. Experiments reversibly perturbing V1 activity and measuring SC responses and behavior would be needed. Moreover, processes associated with the removal of the foveal fixation spot in classic visually-guided saccade tasks need to be understood better. After all, the trigger to generate saccades in our paradigm was a combination of both eccentric target onset and foveal fixation spot removal, but we only studied the neuronal modulations associated with the former but not the latter.

## Resource availability

### Lead contact

Further information and requests for resources and reagents should be directed to and will be fulfilled by the lead contact, Ziad Hafed.

### Materials availability

This study did not generate new unique reagents.

### Data and code availability

- Data: All data reported in this paper will be shared by the lead contact upon request.
- Code: This paper does not report original code.
- Additional information: Any additional information required to reanalyze the data reported in this paper is available from the lead contact upon request.

### Author contributions

YY, TZ, MPB, TM, ZMH collected the data. CT, YY, TZ, MPB, TM, ZMH analyzed behavioral and neuronal data. SP, TM, ZMH processed V1 spike waveform shapes. All authors wrote the manuscript. All authors have read and approved the final version of the manuscript.

## Acknowledgements

We were funded by the Deutsche Forschungsgemeinschaft (DFG; German Research Foundation) through the following projects: SFB 1233 Robust Vision (project number: 276693517); SPP 2205 Evolutionary Optimization of Neuronal Processing (project number: HA 6749/3-2); SPP 2411 Sensing LOOPS: Cortico-subcortical Interactions for Adaptive Sensing (project numbers: 520617944 and 520283985, HA 6749/11-1); BU4031/1-1; BO5681/1-1.

## STAR Methods

### Experimental model and study participant details

#### Animals used

We collected data from SC and V1 neurons in a total of three rhesus macaque monkeys (A, F, and M). A subset of the SC data (from monkeys A and M) were re-analyzed from a previous study ^18^ (particularly, the immediate visually-guided saccade task in that study). Additional SC data were then collected anew, for the purposes of the current project, from monkeys A and F. Data from V1 were also collected anew for the current project from monkeys A and F. The monkeys were all adult (A: 10-13 years, F=14 years, and M=9 years) and male.

#### Animal preparation

The animals, which were group-housed in a dedicated animal facility, were all previously prepared for behavioral and neurophysiological experiments. Briefly, each animal underwent surgeries for head-holder implants, chamber implants (and craniotomies), and scleral search coils. The recording chambers were placed centered on the midline of the skull and tilted backward from vertical by an angle of 38 deg. The chamber centers were aimed, based on structural MRI images, to allow electrodes a straight path towards the SC. In monkeys A and F, we placed the midline chambers (along the anterior-posterior dimension) in such a way to allow targeting dorsal V1 from the posterior halves of these chambers and the whole SC from the anterior halves. For both the SC and V1, we could target both the right and left hemisphere in each animal using the same midline chamber. The scleral search coils allowed us to measure eye movements with very high precision using the magnetic induction technique ^83,84^.

All experiments were approved by the local governmental authorities of the city of Tübingen (Regierungspräsidium Tübingen), and they were in accordance with European Union directives on animal research, along with the German governmental implementations of these directives.

### Method#details

#### Laboratory setup

We used the same experimental setup as that in refs. ^18,49,81^. Briefly, we employed a custom-built modification to PLDAPS ^85^, which interfaced to the PsychToolbox ^86–88^ for display control. Data were stored using the OmniPlex system from Plexon, and the display refresh rate was 85 Hz. The display monitor itself was a CRT display, which was calibrated for luminance and linearized.

In all experiments, the monkeys sat in a dark room, and the CRT display in front of them had a gray background (26.11 cd/m^2^). White stimuli (like the fixation spot; see below) had a luminance of 79.9 cd/m^2^. Also, the distance to the display was 72 cm, giving a display pixel resolution ∼34 pixels/deg.

For the newly recorded data, we used linear multi-electrode arrays (V-Probes from Plexon). We typically employed 16-, 20-, or 24-channel arrays in the SC and 16- or 24-channel arrays in V1. Inter-electrode spacing was 50 μm, except for 2 V1 sessions in which it was 100 μm. For the subset of SC data that were used from the older study ^18^, we used multi-electrode arrays (monkeys A and M) (24-channels). We always sorted neurons offline, as described in more detail below. Thus, all neuronal analyses were performed in the same way for the entire database in this study, whether the data were collected previously or not.

#### Experimental procedures

The behavioral task for the monkeys consisted of an immediate visually-guided saccade task (Fig.1A-D). Each trial started with the appearance of a small white fixation spot (∼10.8 x ∼10.8 min arc dimensions). After approximately 600-1500 ms from fixing gaze on the spot, the spot disappeared and a simultaneous eccentric stimulus (disc of 0.51 deg radius) appeared at a single location across trials per session. The eccentric stimulus was a uniform circular disc of 10%, 20%, 50%, or 100% Weber contrast relative to the gray background (Weber contrast being defined as the absolute difference in luminance between stimulus and background, divided by background luminance). For monkey M (and only sometimes monkey A), we removed the 20% contrast condition, just to reduce the overall number of trials (other tasks had to be performed within the same sessions, requiring some tradeoffs). The stimulus could either be brighter (positive luminance polarity) or darker (negative luminance polarity) than the background, and it was placed at the optimum response field (RF) location of the recorded neurons within a given session. The monkeys’ task was to initiate a foveating eye movement towards the disc as soon as it appeared (with a grace period of 500 ms, but their reaction times were typically significantly faster; see Fig. 1 for examples and Fig. S3C, F, I for each monkey’s results). With their gaze now at the eccentric disc, they were required to maintain fixation on it for another 500 ms before receiving juice or water reward. We typically collected 35-45 trials per condition per session. Figure S14 shows the range of target locations (which were chosen based on the encountered receptive fields) used across all sessions in our experiments.

The above task was always run with other behavioral tasks during the same session. These other tasks were collected for purposes of other scientific questions, but some of them helped us to decide on where to place the eccentric stimulus in a session. Specifically, we ran RF mapping tasks, typically with a delayed, visually- or memory-guided saccade task ^18,49,89^ allowing us to map both visual as well as (if present) motor RF’s. Such RF mapping tasks were run manually, meaning that we manually indicated the desired stimulus location to appear on the next trial. We then changed this location across trials, and we typically covered the entire display (or hemifield in some cases) with a total of 200-300 RF mapping trials. In each RF mapping trial, the monkey fixated a central white or black fixation spot, and a small white spot (the same size as the fixation spot) appeared at the display location that we manually selected for the trial. The evoked visual response on a given electrode channel was connected to an audio speaker to allow us to assess the presence of visual responses on the channel. We visualized and heard visual responses across all channels of a given session, and we could thus assess the average RF location encountered in the session (our electrode tracks were perpendicular to either the SC or V1, meaning that RF’s were generally aligned with each other across the electrode depths). Then, after the initial stimulus onset and evoked neuronal activity, depending on the particular RF mapping task that we were running (e.g. visual or memory-guided saccade), subsequent task events ensued in the trial (e.g. removal of the fixation spot to instruct the monkeys to generate a saccade), but these were less relevant for assessing visual RF locations.

We included a total of 29 SC sessions and 19 V1 sessions in the current study.

### Quantification and statistical analysis

We used the R statistical software for the Generalized Linear Mixed Models (GLMM’s) described below, and we used Matlab for all other analyses.

We detected all microsaccades and saccades using our established methods ^90,91^. We included all trials in which the monkeys were rewarded, regardless of peri-stimulus microsaccades, because we wanted as much variability in visual responses as possible, and we knew that peri-stimulus microsaccades can modulate visual responses for the same stimuli ^19,20,52,64^.

For neuronal analyses, we sorted single units ofline after the experiments using KiloSort 1 and 2, and manually curated in Phy ^92^. We used standard criteria for defining single units. Specifically, the action potential waveforms of a putative single unit had to be self-consistent and contained in well-defined clusters (but some slow, continuous waveform drift was expected and tolerated). Additionally, the waveforms of a single unit had to have fewer than 1.5% violations of a criterion for having a minimal inter-spike interval of 0.8 ms. All neurons included in this study were putative single units, and we did not analyze any multi-units.

We then excluded units that did not exhibit visual bursts. We did this by comparing (via a t-test across trials) firing rates in a 50-ms pre-stimulus interval to those in a visually-driven burst interval defined below, and we then manually inspected all neurons for confirmation. As also mentioned below, note that some neurons responded only for a subset of the tested contrast levels. Thus, in all analyses, we reported the number of neurons that contributed to a given contrast level, which could be smaller than the total number of neurons in our database. In total, our database consisted of 254 SC neurons and 489 V1 neurons.

We detected trial-wise visual response onset latency using a method similar to that used in ref. ^23^. Specifically, we convolved spike times with an asymmetric kernel that did not blur firing rates backwards in time (Fig. S1A). Then, we found the peak firing rate within a trial by searching within a window shortly after stimulus onset (SC two lowest contrasts: 40-110 ms; SC two highest contrasts: 40-100 ms; V1 two lowest contrasts: 30-105 ms; V1 two highest contrasts: 30-95 ms; Fig. S1B). Then, we moved back in time until the trial-wise firing rate dropped below a threshold for at least 5 ms. This threshold was defined based on pre-stimulus activity. In particular, the threshold was defined as 2 standard deviations above the mean pre-stimulus activity across trials (Fig. S1B). The timing of the last spike before crossing this threshold was defined as the trial’s visual response onset latency. In all analyses of trial-by-trial visual response onset latency, we only included a neuron within a given analysis (e.g. 10% dark contrast stimuli) if we could estimate visual response onset latency on at least 60% of all trials within the neuron’s session. This is why neuron numbers reported in Results had different values for different contrasts. For example, neurons with very weak visual responses for low contrast stimuli may not have had sufficient trials with a visual response to warrant inclusion in visual response onset latency analyses.

For visual response strength analyses, we simply counted the number of spikes (per trial) occurring within a short, stimulus-aligned visual response epoch, and we then subtracted the mean number of spikes occurring in a 50-ms baseline epoch across trials (pre-stimulus and ending at stimulus onset). The visual response epoch was area- and stimulus-dependent. Specifically, we typically found that V1 visual bursts were longer lasting than SC visual bursts. Therefore, for V1, our spike count analysis involved counting spikes in an 85 ms time window. Because visual bursts were also contrast-dependent (Fig. 9), the 85 ms window started at 35 ms for the two lowest contrasts (35-120 ms) and at 30 ms for the two highest contrasts (30-115 ms). For the SC, our visual burst epoch was always 60 ms long, and it started at 50 ms for the two lowest contrasts (50-110 ms) and 40 ms for the two highest contrasts (40-100 ms). All of our visual response strength analyses were baseline-subtracted. That is, we always counted spikes in the visual response epoch and subtracted, from the same trial, the average (across trials) number of spikes in a pre-stimulus epoch ending at eccentric stimulus onset. Finally, note that counting spikes within an interval is equivalent to measuring firing rates in the same interval; thus, our approach for visual response strength was not fundamentally different from earlier analyses like those in ref. ^23^.

For pre-stimulus state, we counted spikes, on each trial, within the final 50 ms before stimulus onset. We then correlated this count, on a trial-by-trial basis, to trial-by-trial saccadic reaction time. When a neuron had no pre-stimulus activity (primarily in V1), we designated the neuron as having zero correlation to behavioral variability. For some analyses (e.g. Fig. S11), we estimated baseline firing rate. This was done by convolving our discrete spike times with the asymmetric kernel of Fig. S1A.

Once the neuronal parameters were collected, we took the array of trial-by-trial measures and obtained a Spearman correlation coefficient with the trial-by-trial saccadic reaction times. In some analyses (e.g. Fig. S8), we obtained the Spearman correlation coefficients between two neuronal measures and not between a neuronal measure and behavior. Specifically, we correlated pre-stimulus activity to visual response onset latency. Note that we used Spearman correlations in all analyses because we had no reason to expect a linear relationship between our visual response measures (latencies or strengths) with saccadic reaction times.

We also sometimes assessed variability of neuronal activity independently of correlating to behavior (e.g. Fig. 10). To do so, for each neuron, we measured the standard deviation of a neuronal parameter (e.g. visual response onset latency) across trials within a session. Then, across all neurons, we took the mean and SEM of these standard deviation measures. Thus, the population summaries were estimates of the across-trial standard deviation of a given neuronal parameter. We also did the same to estimate the standard deviation of saccadic reaction time (Fig. S13).

For some analyses, we estimated the cortical depth of V1 neurons. To do so, in each session, we performed current source density (CSD) analysis ^56^, for which we extracted the local field potentials from each electrode channel. These were obtained as described recently ^49,76^. Specifically, we filtered (using a zero-phase-lag filter) the wideband electrode signals to the low frequency band of 0.7-6KHz, and then further low-pass filtered for subsequent analyses ^76^. We then aligned LFP responses (for the highest contrast dark stimulus) to stimulus onset, and took a second spatial derivative across depth, using the method of ^93^. This procedure yielded a CSD profile reflecting the distribution of current sources and sinks across cortical depths. The input layer was defined as the channel exhibiting the strongest current sink (i.e., the most negative CSD value) averaged across the period of 0-100 ms after stimulus onset. Then, after knowing the input channel, we used it and one channel above it or below it (for 50 μm electrode spacings) as the separator between superficial and deep V1 layers. That is, channels above the identified input channel and the one above it were classified as superficial V1 layers; channels below the identified input channel and the one below it were classified as deep V1 layers. For linear electrode arrays with the bigger electrode spacing than 50 μm, we took only the identified channel as the separator between deep and superficial V1. These choices were informed by estimates of cortical thickness in rhesus macaque monkeys ^94^. We had few neurons in the identified input channels (because the channels were the minority in our electrode arrays), so we restricted our analyses to comparisons between deep and superficial cortex. This is justified given how these different cortical sub-units might differentially interact with the SC ^95^.

Also in V1, we analyzed spike waveform shape, to consider whether narrow-spiking or wide-spiking neurons behaved differently from each other. We had a detailed preprocessing pipeline for selecting waveforms. Before estimating the mean waveform shape of each neuron, we first stringently tested whether or not the units were “good”: for each neuron, we took 2000 samples of spike waveforms at the best channel on the probe for that spike, and quantified the distributions of its amplitudes in the pre-spike and spike intervals respectively. Next, we calculated the area under the receiver operating characteristic curve (AUROC) between these distributions to quantify the quality of the spike; the threshold for a “good” neuron was set at an AUROC of at least 0.95. We then ensured that the extracted waveforms for any given neuron did not include accidental noise and were not contaminated by waveforms distorted due to overlaps with waveforms from other neurons. Such contamination would be identifiable as a multimodal amplitude distribution in the spike interval. For such multimodal units, we only accepted waveforms from the largest mode, and finally, summarized it in the mean waveform for that neuron. The waveforms were upsampled from a sampling period of 25 µs to 5 µs using cubic interpolation. After normalizing the waveform amplitudes, we measured the trough-to-peak time interval to assess spike waveform width. Then, inspired by earlier work ^57^, but also driven by our own data distribution, we defined waveforms with a trough-to-peak duration of < 350 μs as being putative interneurons, and waveforms with a longer duration as being putative pyramidal neurons. Note that this value of 350 μs was the threshold for separating the waveform classes; in reality, the average waveform width for the narrow waveforms was significantly smaller (∼200 μs) and consistent with prior studies. We also confirmed that this width threshold was a suitable separator between waveform classes by plotting global histograms of waveform durations in our database, and observing bimodality. In total, we could perform the spike waveform analysis on 198 of our V1 neurons.

For statistics, we found that our correlation coefficient values were generally not normally distributed, by using a Kolmogorov-Smirnov test. Therefore, we employed non-parametric statistical tests throughout our analyses. We had two classes of analyses. In the first, we performed a Mann-Whitney U test comparing correlation values across the conditions of interest. For example, in an analysis like that in Fig. 2A, we tested whether the correlations in the SC were statistically different from those in V1, and we pooled across all contrasts shown in the figure. Then, we compared individual conditions separately (e.g. each contrast level in Fig. 2A). Here, we again used a Mann-Whitney U test but applied Bonferroni correction given the multiple different tested contrasts. In a second class of analyses, we compared correlations to zero. In this case, we used a Wilcoxon Signed-Rank test, and we again applied Bonferroni correction when looking at each contrast level individually. Note that for pre-stimulus activity, and as stated above, when a neuron had no such activity at all (primarily in V1), we designated the neuron as having zero correlation to behavioral variability. As the Wilcoxon Signed-Rank test with reduced sample procedure disregards all samples that equal zero, we reported only the numbers of neurons that excluded this situation in any associated single sample statistical test. Thus, the numbers of neurons indicated in all figures related to pre-stimulus activity are not always the same as the numbers of neurons reported in the associated statistical tests.

For directly comparing visual response latency and visual response strength effects within the same analysis, we also fitted Generalized Linear Mixed Models (GLMM’s). The predicted value in these models was saccadic reaction time, and the fixed effects were visual response latency, visual response strength, and stimulus contrast, all of which were considered continuous and were z-scored prior to the analysis. Neuron identity was included as a random intercept. Saccadic reaction time was modeled using a Gamma distribution with a log link function, since Gamma distributions work especially well for modeling raw reaction time distributions ^96–99^. The model included both main effects as well as interaction terms between visual response latency and contrast, or visual response strength and contrast, respectively. It was specified as:

Saccadic reaction time ∼ visual response latency × stimulus contrast + visual response strength × stimulus contrast + (1|neuron identity)

We created separate GLMM’s for dark or bright luminance polarities. The models were fit using the glmer function from the lme4 package (version 1.1-37) in R ^100^. We used the bobyqa optimizer with a maximum of 2 x 10^5^ function evaluations to ensure convergence. All of our models converged well. The model fit was evaluated using Akaike’s Information Criterion (AIC).

Finally, for the SC neurons, they could be either visual or visual-motor in nature. We assessed this functional dichotomy as follows. We defined a visual window (40-95 after stimulus onset), a pre-motor window (-25-0 ms relative to saccade onset in our main task of Fig. 1), a post-motor window (0-65 relative to saccade onset), and a baseline window (50 ms before stimulus onset). We then calculated the average firing rate for each of the four time windows for each trial, and we performed a multiple comparison Kruskal-Wallis test (α=0.05). If a neuron had a significantly larger activity in the visual window versus the baseline window, it was considered “visual”. If a neuron had a significant difference between the post-motor and baseline window activities and between the pre-motor and post-motor window activities, and if the neuron’s pre-motor activity was lower than its post-motor activity while the pre-motor activity was larger than the baseline activity, the neuron was considered “motor”. These two labels were essentially given out independent of each other. Each neuron that had only the “visual” label was considered “purely visual”, and each neuron that had both labels was considered “visual-motor”.

For additional analyses, we bypassed the categorization into two discrete classes, and instead calculated the visual-motor index (VMI) of each neuron. The VMI was calculated from the delayed, visually-guided saccade task. In this task, we measured visual burst strength as the mean firing rate in the interval 50-100 ms from stimulus onset, and we subtracted the baseline, pre-stimulus firing rate in the interval -100 to -1 ms from stimulus onset. Similarly, we measured the motor burst strength as the peak firing rate +/- 25 ms from saccade onset, and we subtracted a baseline rate obtained as the average firing rate in the final 100 ms before the end of the delay period before the instruction to trigger the saccade (-100 to -1 ms). The VMI was calculated as the visual burst measure minus the motor burst measure, divided by the sum of the two, and it could thus cover a range of values from +1 to -1. If a neuron was significantly suppressed in the motor epoch, its VMI value was assigned to +1. After calculating VMI’s, we binned them into bins of +/- 0.1, with a step size of 0.05, and we then plotted the average correlation coefficient value of all neurons falling within a given VMI bin.

All results of statistical tests, including numbers of observations, are listed in the text of Results and/or the figure legends.

## Supplemental information

Document S1. Figures S1-S14.

---

**Figure S1.**
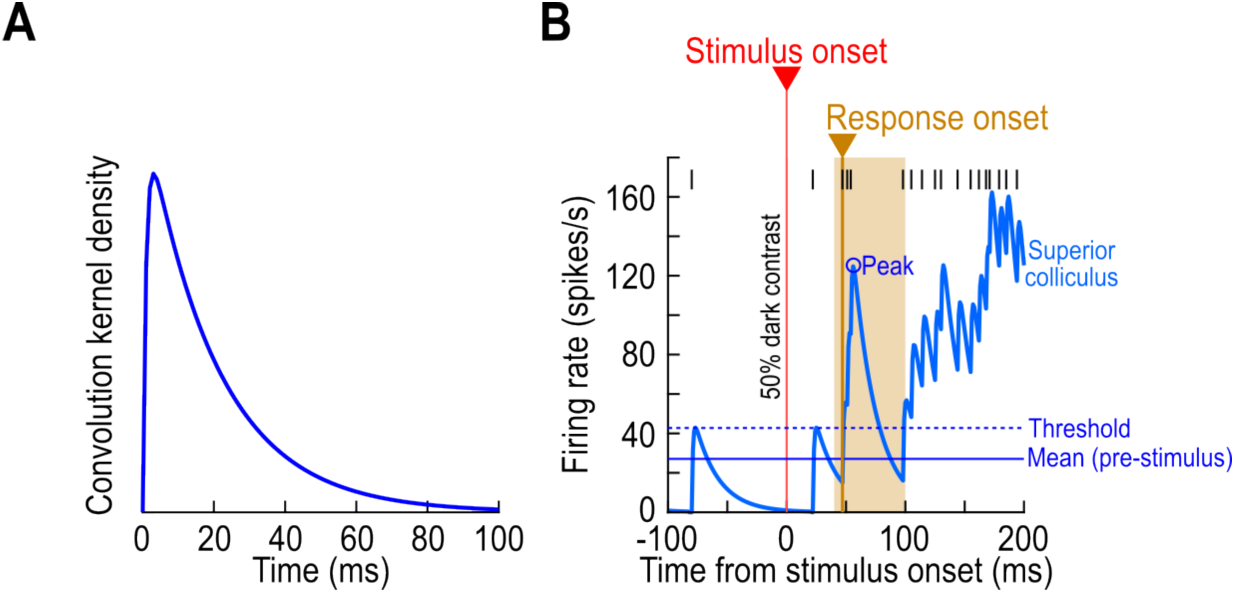
Estimating trial-wise visual response onset latency from a neuron’s spike times, related to Fig. 1. **(A)** The convolution kernel that we employed in order to estimate firing rates from individual-trial spike times. Each spike in a trial was convolved with this kernel density function. **(B)** Example trial from an SC neuron illustrating our method for estimating trial-wise visual response onset latency. The black vertical tick marks in the top of the figure are the individual spike times emitted by the recorded neuron on the shown trial. The blue curve is the estimated firing rate curve of the trial, given the convolution of the spike times with the kernel in **A**. We estimated visual response onset latency by comparing to pre-stimulus activity. Specifically, after finding the peak firing rate (circle labeled “Peak”) within a search window (brown background rectangle), we moved backwards in time until the firing rate dropped below a threshold. The threshold was defined based on measurements of pre-stimulus activity (Methods).

**Figure S2.**
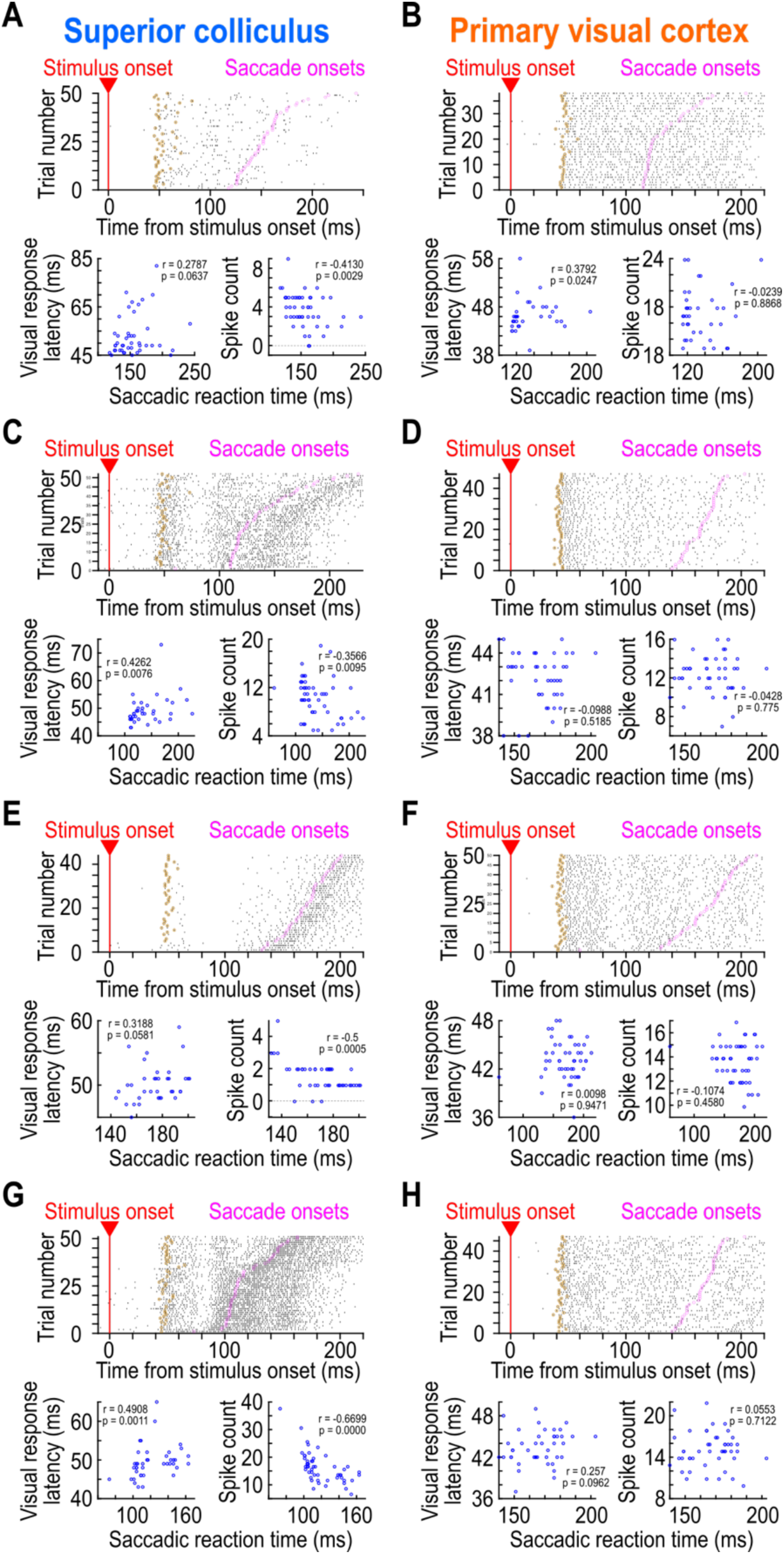
Additional example neurons from both brain areas, exhibiting results that are representative of the whole population, related to Fig. 1. This figure is formatted identically to Fig. 1, and it shows four more example SC neurons (**A**, **C**, **E**, **G**) and four more example V1 neurons (**B**, **D**, **F**, **H**). Note how the first example SC neuron (**A**) was a purely visual neuron, because it did not emit a saccade-related motor burst at the time of saccade onset. Also see Fig. 3 for the firing rate estimates of the neurons.

**Figure S3.**
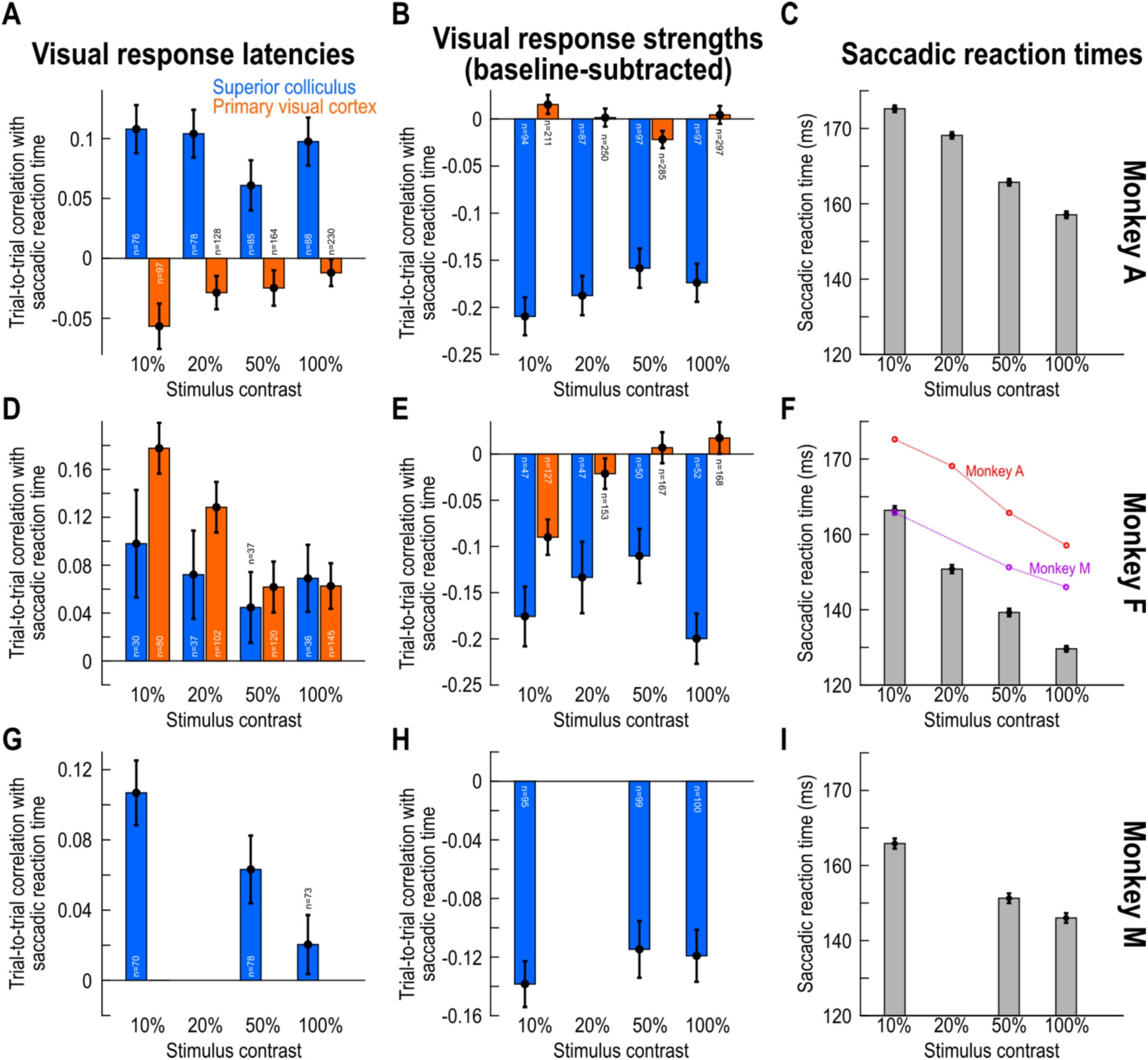
Individual monkey results, related to Fig. 2. **(A, B)** Similar to Fig. 2A, B, but now for only the data obtained from monkey A. Note how the results, especially for the SC, were consistent with those of Fig. 2. **(C)** The same monkey’s saccadic reaction times as a function of stimulus contrast (dark stimuli shown; similar trends could be observed for bright stimuli, save for the fact that saccadic reaction times were generally slightly faster for darks than brights). Each bar shows the average saccadic reaction time across all trials from all sessions in this monkey, for a given stimulus contrast, and the error bars denote SEM. **(D-F)** Similar analyses but now for monkey F (the y-axis ranges are tailored for each monkey’s results). Note how the correlation coefficients for V1 in **D** were higher than in monkey A (compare the V1 data in this panel to those in **A**). Interestingly, monkey F had much faster saccadic reaction times than the two other monkeys; for example, in **F**, we superimposed the monkey A and M saccadic reaction times of **C**, **I** in red and purple color, respectively, for easier comparison. This suggests that for more reflexive monkeys like monkey F, sensory drive from V1 (**D**) might matter more for saccadic reaction times than in other cases. In terms of the SC (blue bars), this monkey exhibited consistent results with monkey A. **(G-I)** This consistency was also clearly evident in monkey M. Note that in this monkey, we did not test 20% contrast levels (Methods). Thus, across all monkeys the SC results were highly consistent. All other conventions are similar to those in Fig. 2. Also see Fig. S5 for further classification of the SC neuron types.

**Figure S4.**
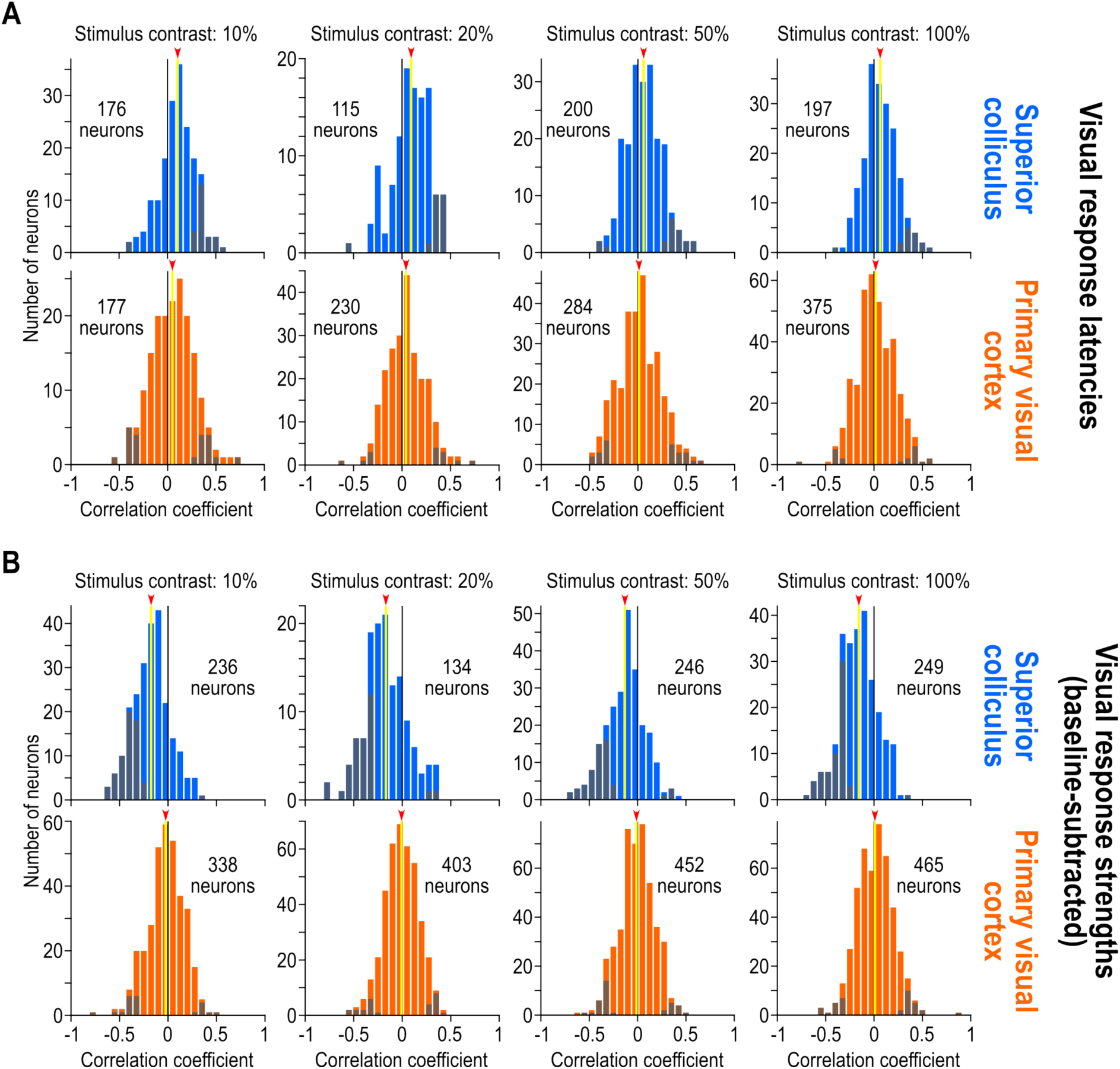
Raw data underlying Fig. 2, related to Fig. 2. **(A)** Raw histograms showing the underlying raw data of Fig. 2A. Each plot shows the results from one contrast level. The x-axis in each plot indicates the Spearman correlation coefficient value, and the y-axis shows the number of neurons exhibiting this correlation coefficient value. The yellow vertical line in each plot (and the red arrowhead on its top) shows the mean of the shown population, and the dark bars indicate the neurons that individually had significant Spearman correlation coefficient values. Note how the SC population had consistently positive correlation coefficients, whereas the V1 population had a mean correlation coefficient very close to zero. **(B)** Raw histograms showing the underlying raw data of Fig. 2B. Note how the SC results were even stronger than in **A**, but in the negative direction, consistent with Fig. 2. Moreover, the V1 distributions were all centered around zero, again consistent with Fig. 2. Also see Fig. S6 for further delineation of the SC data as a function of whether the neurons were visual or visual-motor neurons.

**Figure S5.**
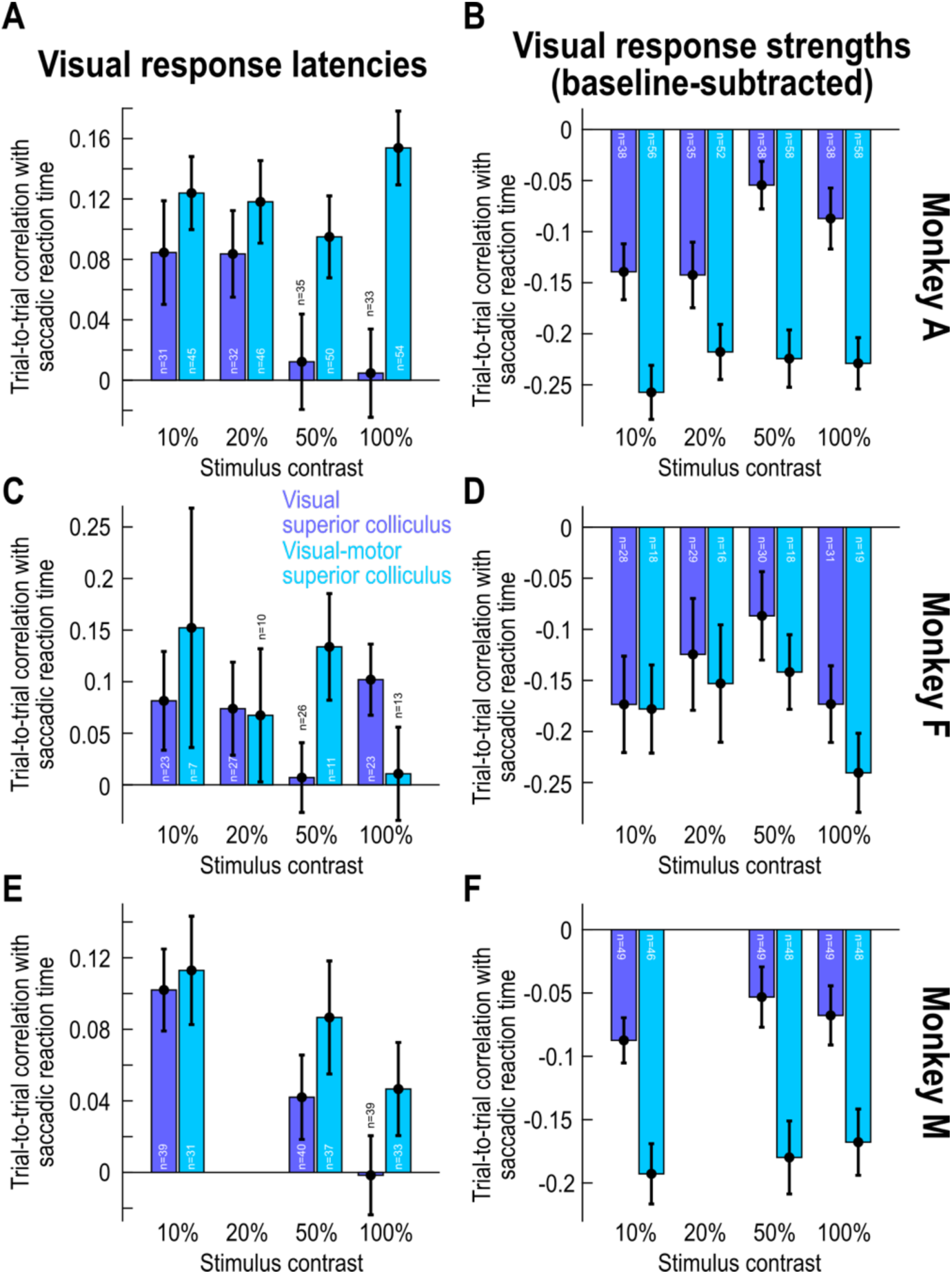
Individual monkey results for Fig. 4, related to Fig. 4. This figure is formatted similarly to Fig. S3 above, but now separating the SC visual and visual-motor neurons. All monkeys showed consistent results. The figure is formatted identically to Fig. 4, except that we do not show the V1 data, which are shown in Fig. S3.

**Figure S6.**
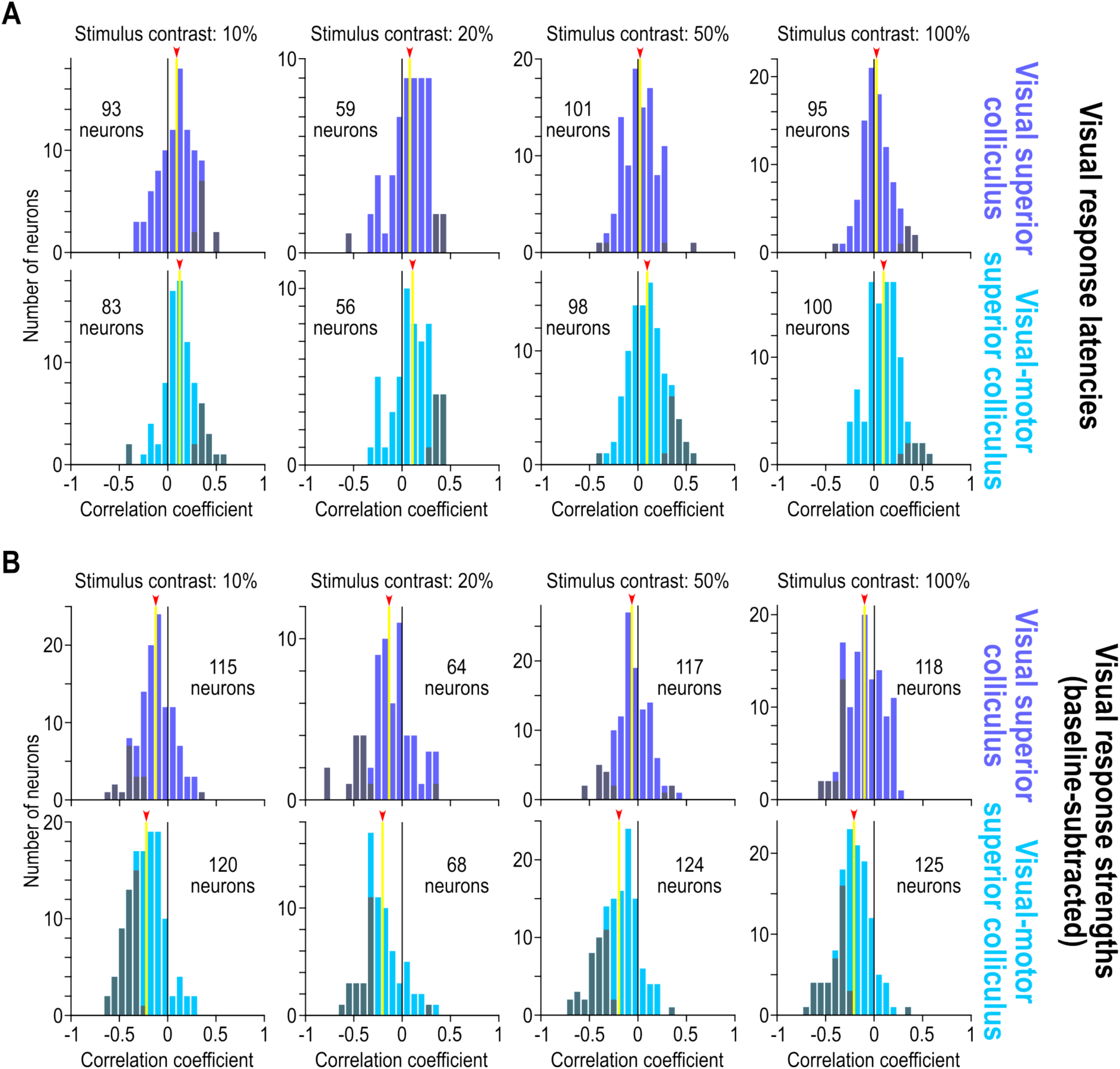
Raw data underlying Fig. 4, related to Fig. 4. **(A)** Similar to Fig. S4A, but now comparing SC visual (top row) to SC visual-motor (bottom row) neurons. Note how both the visual and visual-motor neurons showed positive correlation coefficient values across the population (yellow vertical lines indicating the means), and also note how the effects were stronger for the visual-motor neurons (compare the yellow lines), as summarized in Fig. 4A. **(B)** The effects were even stronger for visual response strengths in the SC, but in the negative direction, again as summarized in Fig. 4B. All other conventions are similar to those in Fig. S4.

**Figure S7.**
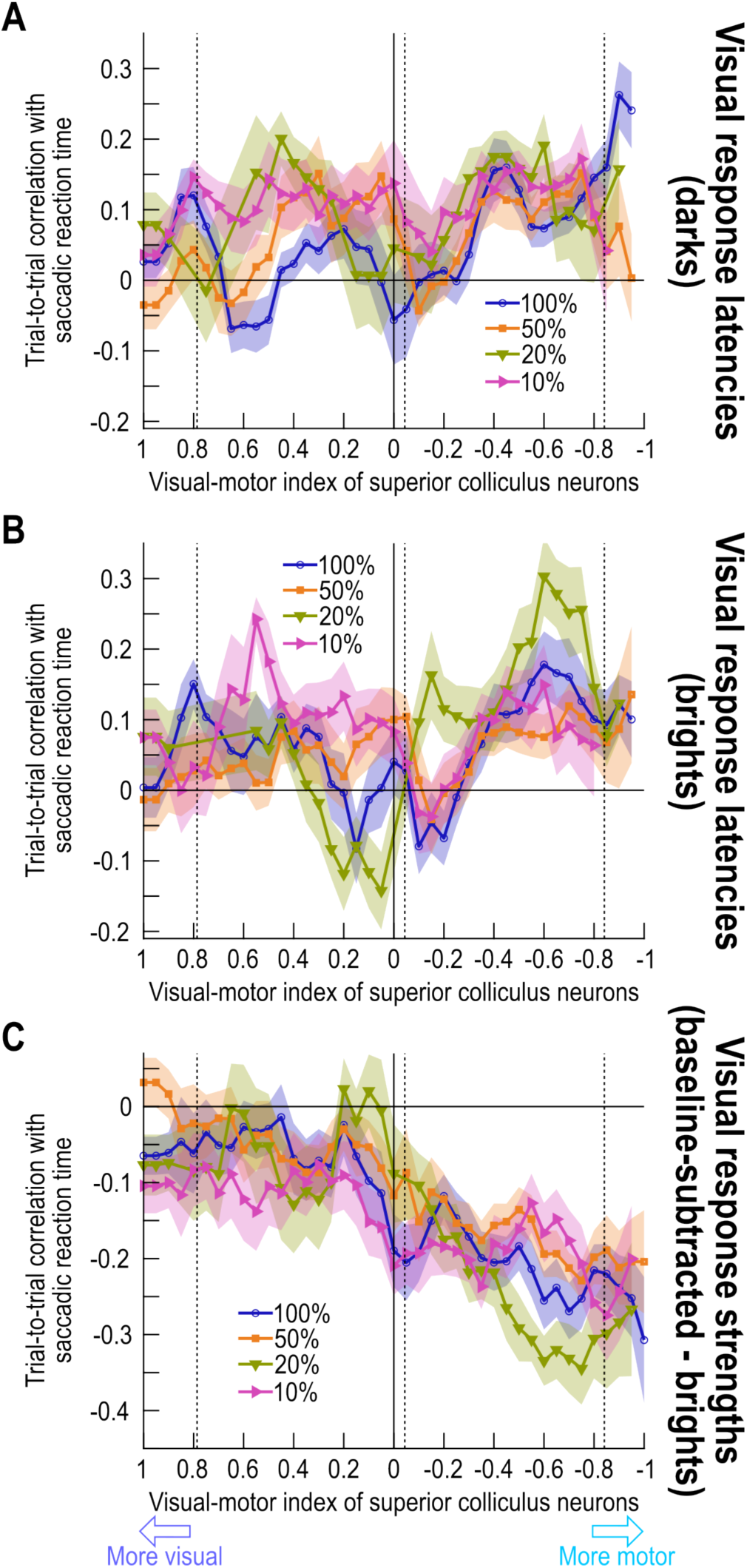
Dependence of Spearman correlation coefficient values on functional depth within the SC, as assessed via visual-motor indices, related to Fig. 5. **(A)** Similar to Fig. 5, but now plotting the Spearman correlation coefficients (relating SC visual response latency, as opposed to visual response strength in Fig. 5, to saccadic reaction time) as a function of the visual-motor index (VMI) of the recorded SC neurons. More visual VMI’s (left side of the x-axis) are associated with more superficial SC neurons, and more negative VMI’s (right side of the x-axis) are associated with more deep SC neurons. The correlation coefficients increased with increasing functional depth within the SC, consistent with the results of Fig. 4. Note that the increase with the shift of VMI’s towards more motor values (rightward on the x-axis) was generally weaker than the effects that we observed with visual response strengths in Fig. 5. This is consistent with our earlier observations that visual response strength, not latency, was the better predictor of saccadic reaction times in the SC data (Figs. 2, 4). **(B)** Same as **A** but for bright visual stimuli. Similar trends were observed. **(C)** Same as Fig. 5 but for bright stimuli. Similar trends were observed. Error bars denote SEM.

**Figure S8.**
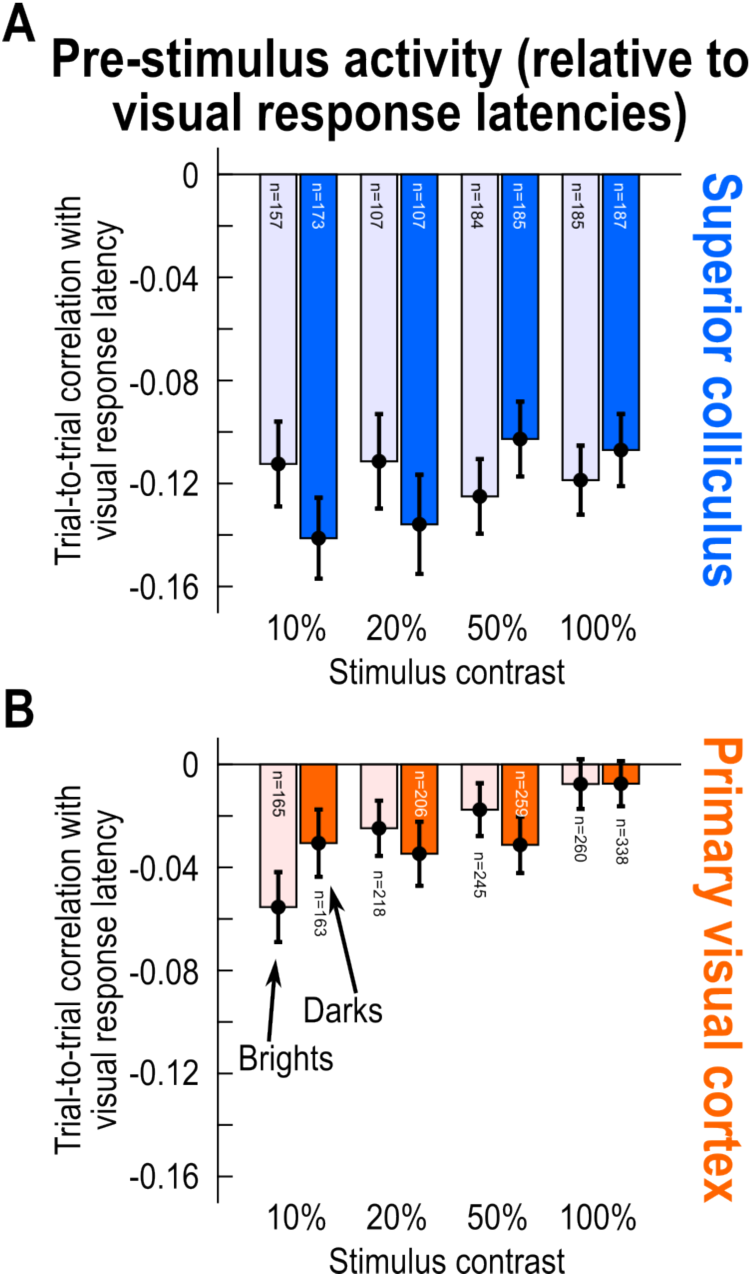
Relating trial-by-trial variability in pre-stimulus (baseline) activity to trial-by-trial variability in visual response latency, in both the SC and V1, related to Fig. 8. **(A)** For each SC neuron and stimulus condition, we measured trial-by-trial pre-stimulus activity and correlated it with trial-by-trial visual response onset latency. Negative correlations mean weaker pre-stimulus activity for later stimulus-evoked visual response onset latency. The results indicate that visual response onset latencies were a function of pre-stimulus SC state, and not completely independent of it. More importantly, there was no difference in the effects between dark and bright contrasts (U=418557, z=0.14, p=0.8885, n_brights_=1067, n_darks_=689; Mann-Whitney U test comparing darks and brights across contrasts). This suggests that the weaker correlations to behavior (rather than visual response onset latency) in pre-stimulus activity for bright contrasts (Fig. 8C) was mediated by a mechanism other than a link between pre-stimulus activity and visual response latency. Potentially, this other mechanism could include visual response strength correlations (Figs. 8B, S9) because our visual response strength analyses were always independent of pre-stimulus activity (Methods). **(B)** For V1, the visual response latencies for low contrast stimuli (except for 20% brights) did generally depend on pre-stimulus activity (10% brights: W=2962, z=-3.90, p<0.0001, n=146; 10% darks: W=3547, z=-2.41, p=0.0158, n=150; 20% brights: W=7040, z=-1.92, p=0.0548, n=196; 20% darks: W=5295, z=-2.64, p=0.0084, n=190; Wilcoxon signed-rank test against zero). This could partially explain why V1 visual response latencies were better correlated to saccade timing with low contrast bright stimuli (Fig. 8D). Error bars denote SEM, and the numbers of neurons included in each analysis are included in the figure.

**Figure S9.**
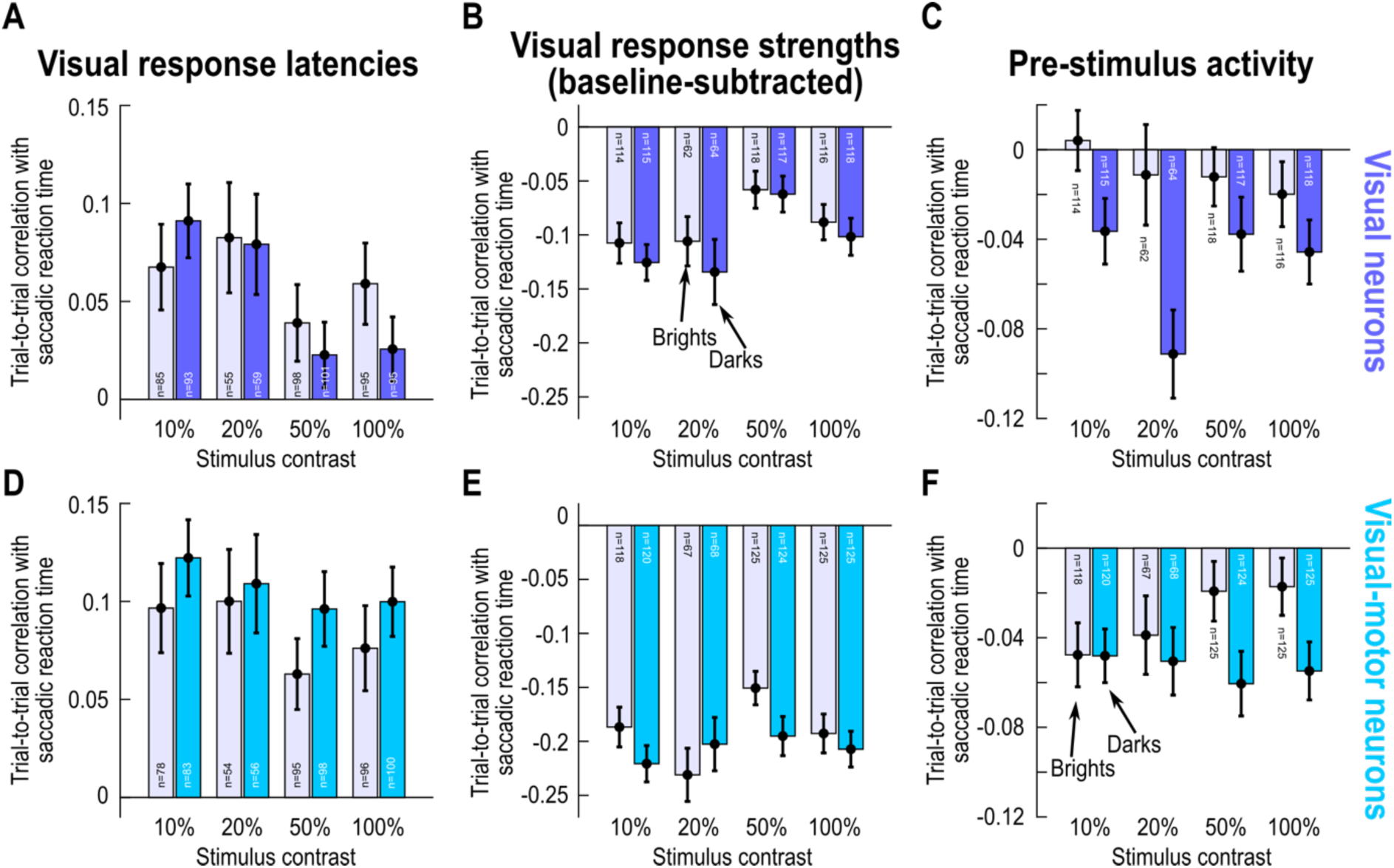
Similar dependencies on dark versus bright contrasts in both SC visual and visual-motor neurons, related to Fig. 8. **(A-C)** Same analyses as in Fig. 8A-C but only for the visual-only neurons of our SC population. Saturated bars show results for dark contrasts, and faint bars show results for bright contrasts. In terms of visual response onset latency (**A**), there was no statistically significant difference between dark and bright contrasts (U=116511, z=-0.84, p=0.4007, n_brights_=333, n_darks_=348, Mann-Whitney U test comparing brights and darks across contrast levels). This was also true for visual response strength (**B**) (U=168217, z=-0.75, p=0.4541, n_brights_=410, n_darks_=414; Mann-Whitney U test comparing brights and darks across contrast levels). However, for pre-stimulus activity (**C**), there were weaker correlations with trial-by-trial saccadic reaction times in the case of bright versus dark stimuli (U=159130, z=-3.41, p=0.0006, n_brights_=410, n_darks_=414), but only 20% contrast was individually significant after Bonferroni correction. **(D-F)** Same analyses as in Fig. 8D-F but only for the visual-motor SC neurons of our population. For visual response strength (**E**) (U=183437, z=-1.97, p=0.0493, n_brights_=435, n_darks_=437) and pre-stimulus activity (**F**) (U=181828, z=-2.40, p=0.0163, n_brights_=435, n_darks_=437), there were weaker effects in the case of bright contrasts; however, none of the individual contrast comparisons reached significance after Bonferroni correction. Thus, the generally weaker effects for bright contrasts in Fig. 8 were consistent across SC neuron types.

**Figure S10.**
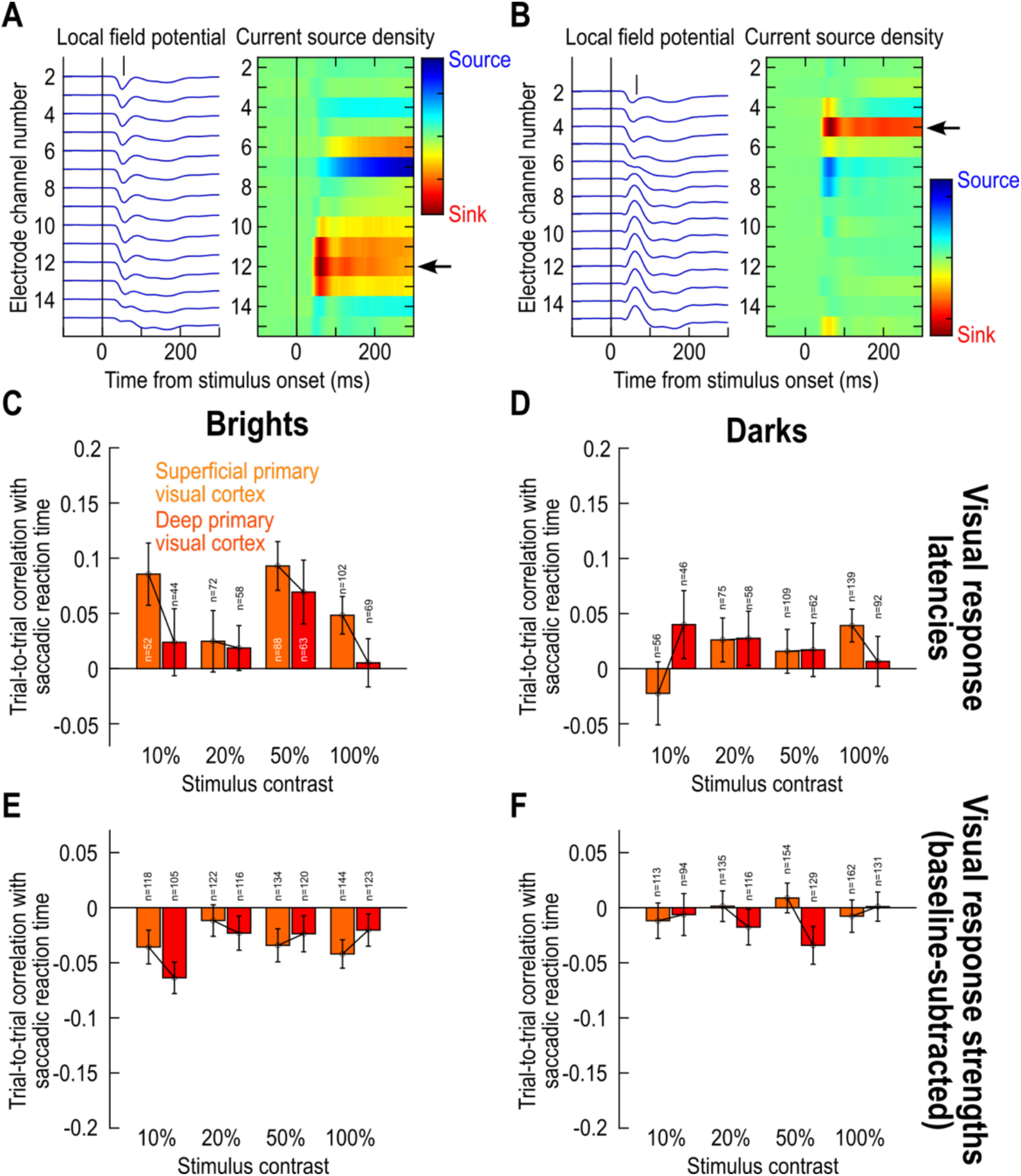
No consistent difference in effects between superficial and deep V1 layers, related to Fig. 8. **(A)** Using current source density (CSD) analysis (Methods), we classified our V1 neurons according to whether they were in putative superficial or deep V1 layers. In the left panel, we plotted stimulus-evoked local field potential (LFP) deflections as a function of time from stimulus onset; each trace shows the deflection from one channel of the recording electrode array from one example session. In the right panel, we used the CSD pattern inferred from the LFP deflections of the left panel, and we identified the sink at channel 12 (highlighted with a black arrow); note how this was also the channel with the earliest stimulus-evoked responses. Thus, for this session, channel 12 corresponded to the putative input layer: we classified neurons above this channel as superficial V1 neurons, and neurons below as deep V1 neurons (Methods). **(B)** In another example session, the putative input layer was identified at channel 5 of the electrode array. Thus, neurons from channels above channel 5 were the superficial neurons, and neurons from channels below were the deeper neurons. **(C-F)** Across all sessions, after alignments like in **A**, **B**, we repeated our analyses but only for subsets of V1 neurons. There were no systematic differences in effects as a function of V1 depth. Only in the case of bright stimuli (**C**) were there larger correlations in the superficial neurons (U=89797, z=1.97, p=0.0494, n_superficial_=314, n_deep_=234; Mann-Whitney U test across contrasts). Note that we did not analyze effects related to pre-stimulus activity as a function of V1 depth because such activity was very low anyway, and also because it did not strongly depend on such depth (see Fig. S11).

**Figure S11.**
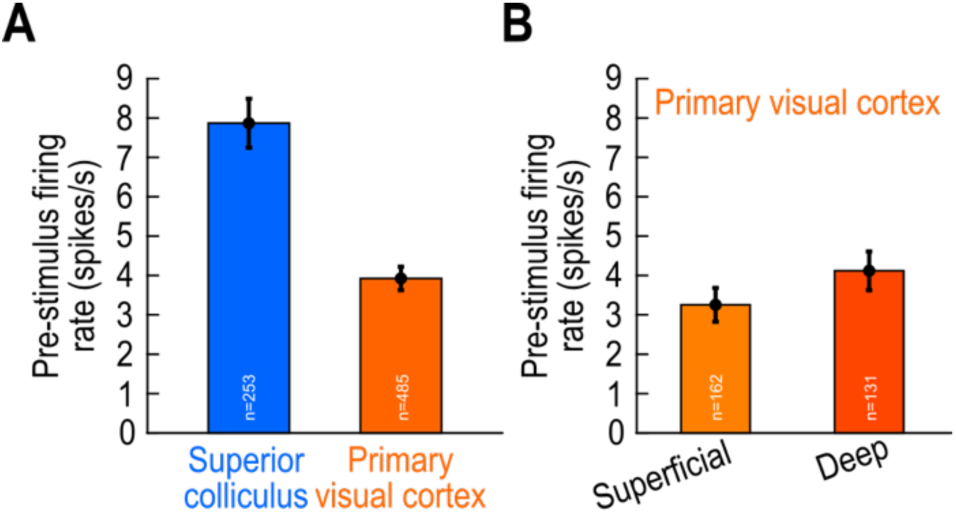
Weaker pre-stimulus (baseline) activity in V1 than in the SC, related to Fig. 8. **(A)** SC neurons possessed twice the pre-stimulus (baseline) firing rate of V1 neurons. **(B)** Neurons in the superficial and deep V1 layers had similar pre-stimulus firing rates. The input layers of V1 had even weaker pre-stimulus activity (1.44 spikes/s +/- 0.395 spikes/s SEM; n=38 neurons). Error bars denote SEM, and the numbers of neurons in each analysis are shown in the figure.

**Figure S12.**
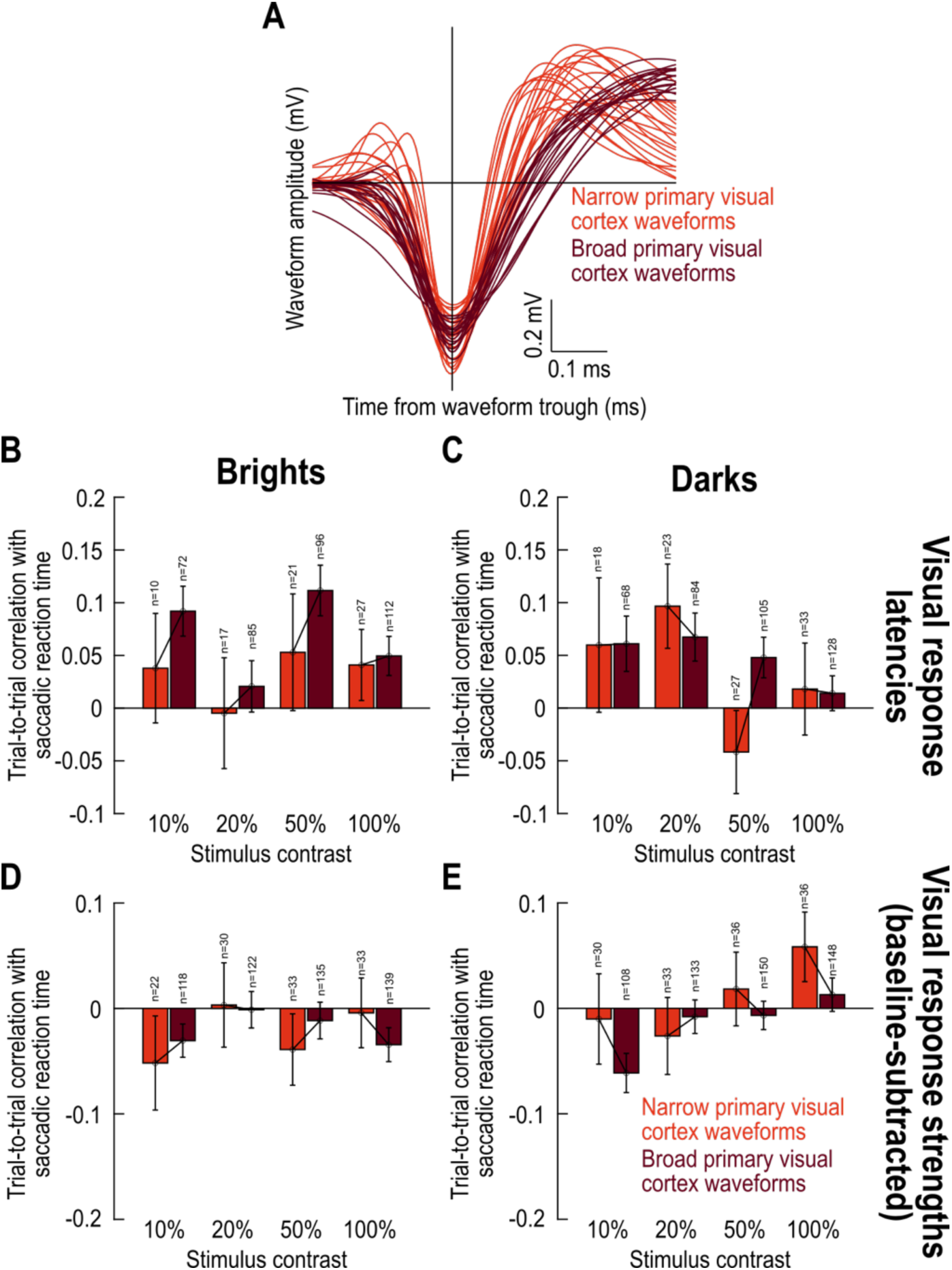
No consistent difference in effects between narrow- and broad-spiking V1 neurons, related to Fig. 8. **(A)** Twenty randomly chosen neurons having a broad action potential waveform (dark color), and twenty having a narrow action potential waveform (lighter color; Methods). **(B-E)** None of our analyses revealed any significant differences between narrow- and wide-spiking neurons in V1, in terms of correlations to behavioral variability.

**Figure S13.**
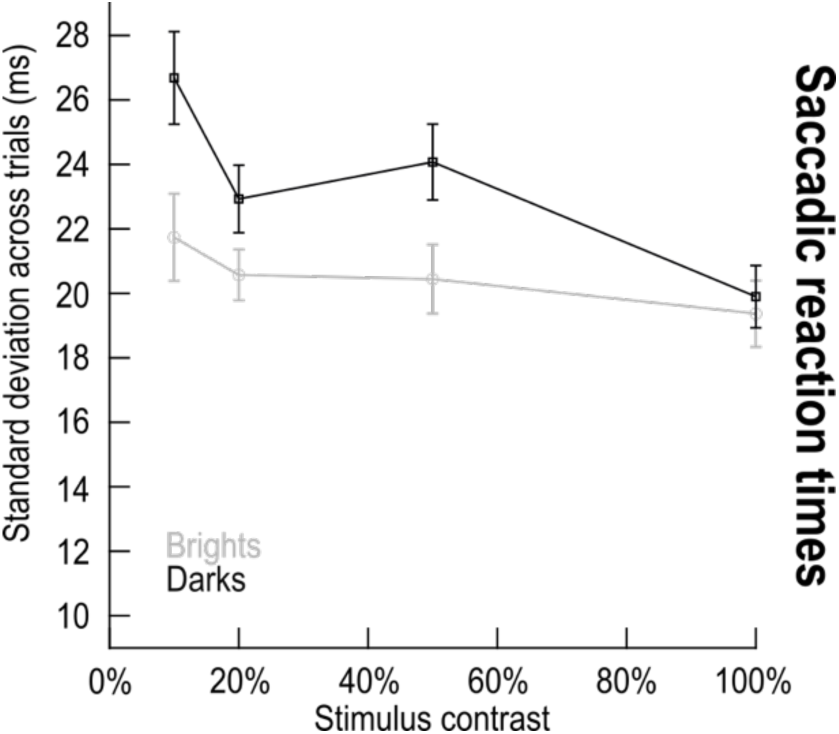
High variability in foveation timing, related to Fig. 10. For each behavioral session, we measured the standard deviation of saccadic reaction times across trials. The shown summaries are averages and SEM ranges across sessions (dark and square symbols indicate dark contrasts, and bright and circle symbols indicate bright contrasts). In comparison to Fig. 10, the timing variability in behavior was much higher than the timing behavior of visual response onset latencies in both the SC and V1.

**Figure S14.**
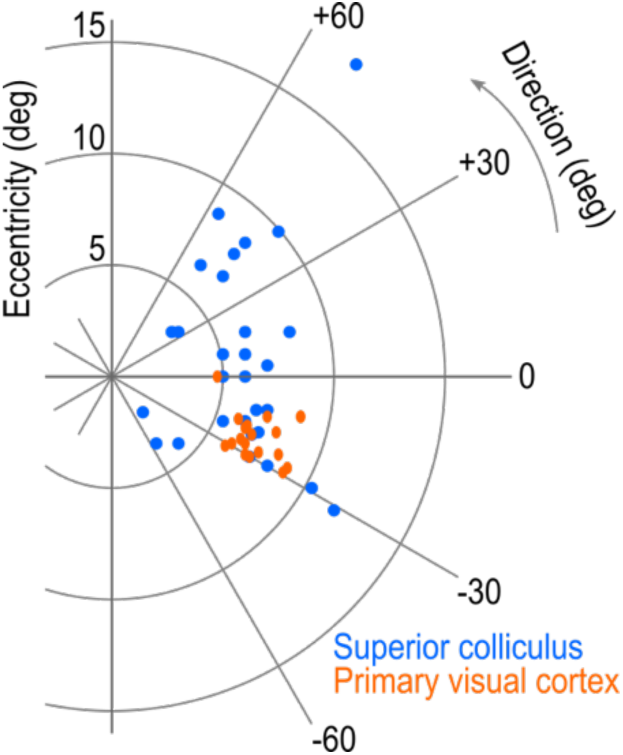
Sampled eccentricities and direction in SC and V1, related to Fig. 2. This figure shows the saccade target locations used across sessions in the SC (blue circles) and V1 (orange ovals). Such locations were chosen according to the average receptive field location in each electrode penetration (Methods). In both brain areas, we had similar ranges of eccentricities of the saccade targets. In the SC, we sampled both upper and lower visual field stimulus locations.

